# KIF18A Inhibition as a Therapeutic Strategy in Cancers with Rb Pathway Inactivation

**DOI:** 10.64898/2026.04.14.718587

**Authors:** Celia Andreu, Arshi Arora, Song Chen, Chaithanya Vedula, Adriana Roopnariane, Sarah Bettigole, Nazario Bosco, Aamir Dohadwala, the SOVI-2302 Investigators, the VLS-1488-2201 Investigators, David P. Southwell, Neil J. Ganem, Timothy Bowler, Samuel F. Bakhoum

## Abstract

KIF18A inhibition has emerged as a therapeutic strategy for chromosomally unstable cancers, but clinical development is limited by the absence of a deployable predictive biomarker. Here we identify strong, diffuse p16INK4a expression, a well-established surrogate marker of Rb-pathway inactivation, as a predictive biomarker of response to KIF18A inhibition, and show that Rb-pathway inactivation marks a biologically distinct subset of cancers sensitive to this therapeutic approach. In sensitive models, low Rb activity is associated with robust spindle assembly checkpoint signaling and prolonged mitotic arrest following KIF18A inhibition. Weakening the spindle assembly checkpoint in this context is sufficient to confer resistance. Across three independent pan-cancer sensitivity datasets generated with distinct KIF18A inhibitors, Rb-pathway altered models were significantly more sensitive than histology-matched Rb-intact comparators, with the strongest association observed in cancers harboring direct RB1 loss or inactivating mutation. Guided by this mechanism, we retrospectively analyzed p16INK4a expression by immunohistochemistry (IHC) in pre-treatment tumor biopsies from 79 heavily pre-treated high-grade serous ovarian cancer patients across three dose-escalation or expansion cohorts and treated with two different KIF18A inhibitors (sovilnesib and VLS-1488) sharing a common mechanism of action. p16INK4a-high tumors showed substantially higher objective response rates than their p16INK4a-low counterparts (36.0% versus 2.2%; P = 0.0002) and markedly longer progression-free survival (median 24.3 versus 7.9 weeks; hazard ratio, 0.16; P < 0.0001). These findings establish p16INK4a as a mechanistically-based, clinically implementable biomarker of clinical response to KIF18A inhibition that is poised to support pan-cancer development of KIF18A inhibitors guided by Rb-pathway inactivation.

## INTRODUCTION

Chromosomal instability, defined by persistent mis-segregation of whole chromosomes or chromosome fragments during cell division, is a pervasive feature of human cancer^1^. CIN contributes to intratumoral heterogeneity, therapeutic resistance, metastatic progression, and adverse clinical outcome across multiple tumor types^2–4^. Despite its prevalence, CIN has remained difficult to exploit therapeutically because most agents that perturb mitosis also injure normal proliferating tissues, particularly bone marrow and gastrointestinal epithelium. A central challenge, therefore, has been to identify mitotic dependencies that arise from the chromosomally unstable state itself rather than from proliferation alone.

KIF18A has emerged as an attractive candidate in this setting^5–9^. KIF18A is a plus end directed mitotic kinesin that regulates chromosome congression and progression through mitosis^10–12^. It has been shown independently that cancer cells with CIN^5^, aneuploidy^8^, or whole genome doubling^9^ exhibit selective dependence on KIF18A, whereas near diploid cells generally do not. This biology suggests a therapeutic window distinct from that of conventional antimitotic therapy. Small molecule KIF18A inhibitors have shown minimal effects on human bone marrow mononuclear cells and foreskin fibroblasts, and inhibition of DNA synthesis in proliferating mammary epithelial cells and activated human T lymphocytes only at concentrations well above those required in sensitive cancer cell lines^6,7^. In sensory neuron assays, KIF18A inhibitors had no meaningful effect on neurite outgrowth except at high micromolar exposure. Together, these data support the premise that KIF18A inhibition targets a cancer specific mitotic state while comparatively sparing normal somatic cells.

In the first-in-human phase I/II study of VLS-1488, a highly potent and selective KIF18A inhibitor, 52 patients with advanced solid tumors were treated across dose levels from 50 to 800 mg once daily^13^. No dose limiting toxicities were observed and the maximum tolerated dose was not reached. Fewer than 45% of patients experienced treatment-related adverse events of any grade, fewer than 16% experienced grade 3 treatment-related adverse events, and no patient experienced a treatment-related event above grade 3. Among 20 patients with advanced high grade serous ovarian cancer, most of whom were platinum resistant and had received a median of five prior lines of therapy, 7 of 17 response-evaluable patients demonstrated tumor reduction, including 3 partial responses. These early findings are noteworthy in a heavily pretreated platinum resistant population, demonstrating tolerability along with efficacy.

A critical limitation for further clinical development of KIF18A inhibitors is the lack of a predictive biomarker. Initial genetic and pharmacologic studies suggested that whole genome doubling (WGD), aneuploidy, fraction genome altered (FGA), and broader measures of CIN may enrich for KIF18A dependency. However, these descriptors define broad biological states rather than a clinically feasible patient selection strategy. Chromosomal and karyotypic abnormalities have been deemed necessary for robust sensitivity to KIF18A inhibition, but they may not be sufficient, implying that only a biologically distinct subset of tumors with chromosomal abnormalities is truly dependent on this target.

To identify the subset of chromosomally unstable tumors most likely to respond to KIF18A inhibition, we focused on the spindle assembly checkpoint, which is required for KIF18A inhibitor response^14^. One of the key upstream regulators of spindle assembly check-point signaling is the Rb pathway, acting through the E2F family of transcription factors. E2F controls transcription of several mitotic and checkpoint regulators, including MAD2^15^, which led us to investigate Rb-pathway inactivation and p16INK4a, a well-established marker of that state^16,17^. Here we show that Rb-pathway inactivation defines a biologically distinct subset of chromosomally unstable tumors that is selectively vulnerable to KIF18A inhibition, and that strong diffuse p16INK4a expression provides a clinically deployable immunohistochemical readout of this state. In a retrospective clinical analysis of pre-treatment tumor samples from heavily pretreated high-grade serous ovarian cancer (HGSOC), this association was observed across two KIF18A inhibitors that share the same mechanism of action, VLS-1488^7^ and sovilnesib^6^, supporting an on-mechanism rather than compound-specific effect. Together, these findings provide a mechanistically based framework for prospective biomarker-enriched development of KIF18A inhibitors.

## RESULTS

### KIF18A inhibitor sensitivity is associated with prolonged mitotic arrest and elevated Mad1

To identify the biological state that underlies response to KIF18A inhibition, we began with two complementary cell-line cohorts: a panel comprising primarily esophageal cancer-derived cell lines along with a normal esophageal mucosal cell line (HET1A), a non-dysplastic Barrett’s esophageal cell line (CP-A), and a triple-negative breast cancer cell line (HCC1806) was used to define the dynamic mitotic phenotype of response at single-cell resolution through live-cell imaging, and an ovarian cancer panel, prioritized for biomarker discovery since the focus of our clinical development is high-grade serous ovarian cancer (HGSOC). Across the esophageal panel, KIF18A inhibitor treatment separated sensitive and insensitive models on the basis of growth inhibition (**Fig. 1a**). Live-cell imaging showed that this difference reflected a marked divergence in mitotic behavior: whereas most cell lines exhibited some form of mitotic delay upon treatment with KIF18A inhibitors, this delay was much more pronounced in the sensitive cell lines, whereas insensitive lines ultimately progressed through mitosis despite treatment (**Fig. 1b**). Cell-fate mapping further showed that sensitive cells frequently died during mitosis, died after mitotic slippage, or exited mitosis with abnormal outcomes including micronucleation, whereas insensitive cells more often completed apparently viable divisions (**Fig. 1c**). Importantly, baseline mitotic duration did not correlate with drug sensitivity (**Supplementary Fig. 1**), indicating that response was not simply a consequence of intrinsically longer mitosis. These findings suggest that cell lines with robust spindle assembly checkpoint signaling were more likely to exhibit sensitivity to KIF18A inhibition.

**Fig. 1.**
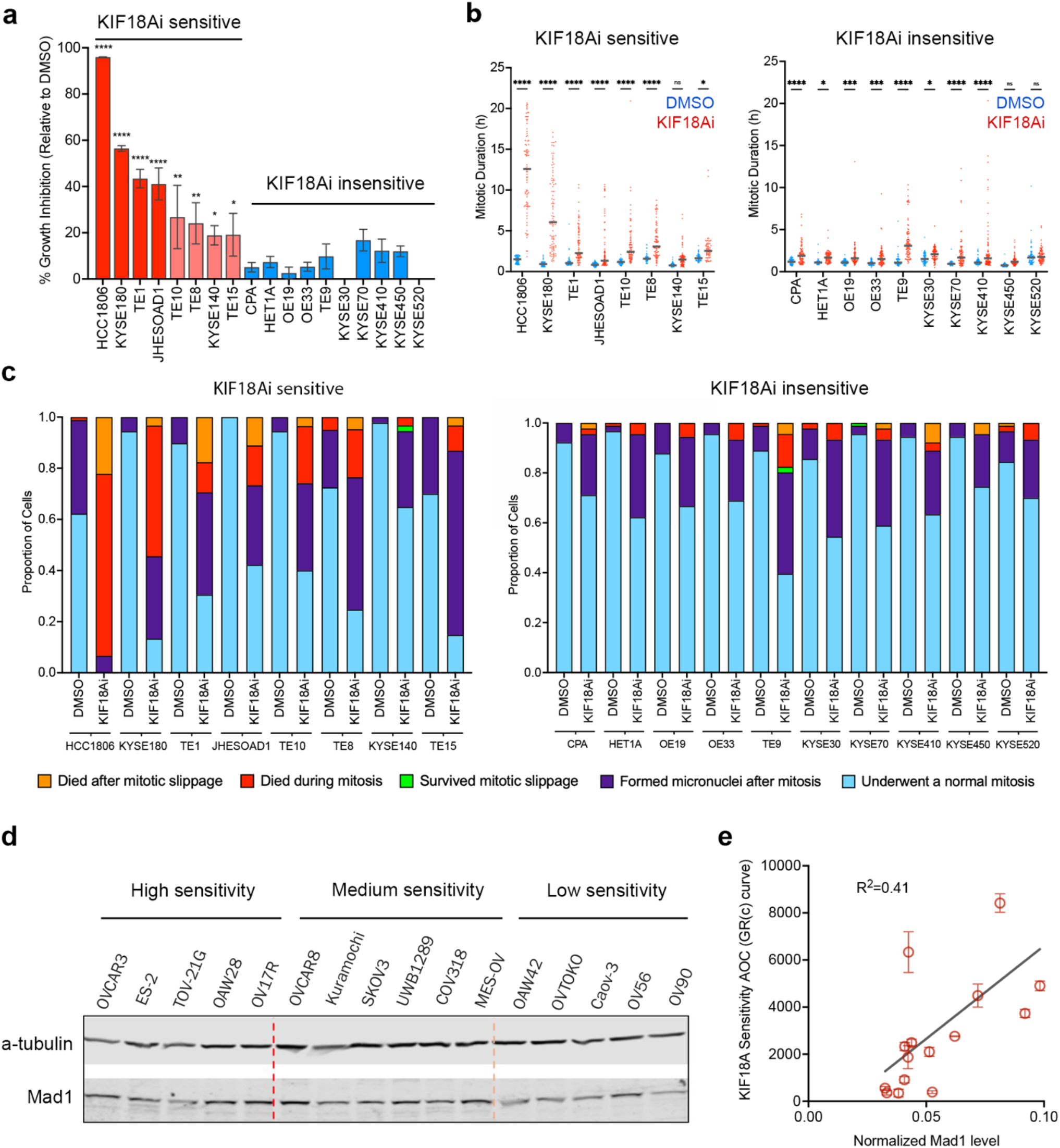
KIF18A inhibitor sensitivity is associated with prolonged mitotic arrest and elevated Mad1. **a**, Growth inhibition after KIF18A inhibitor (VLS-1272) treatment across a panel of cell lines, grouped by sensitivity and shown relative to DMSO. **b**, Mitotic duration – obtained from live cell imaging – in KIF18A inhibitor-sensitive and KIF18A inhibitor-insensitive cell lines following treatment with DMSO or VLS-1272. Each data point represents one cell. **c**, Cell fate mapping for representative KIF18A inhibitor-sensitive and KIF18A inhibitor-insensitive esophageal cancer cell lines treated with DMSO or KIF18A inhibitor. Colors denote the indicated mitotic outcomes. **d**, Immunoblot analysis of Mad1 across an ovarian cancer cell line panel grouped by KIF18A inhibitor sensitivity. α-tubulin served as a loading control. **e**, Correlation between normalized Mad1 protein abundance and KIF18A sensitivity score across the ovarian cancer cell line panel.

We then asked whether the expression of key spindle assembly checkpoint signaling components was associated with response to KIF18A inhibition, using an ovarian cancer cell line panel that would serve as the basis for biomarker discovery. This panel exhibited varying sensitivity to KIF18A inhibitors (**Supplementary Fig. 2**). Indeed, the abundance of Mad1, an essential spindle assembly checkpoint protein^18^ and a binding partner of Mad2 and essential for the latter’s recruitment to the kinetochore, was higher in the more sensitive ovarian lines and correlated with the KIF18A sensitivity score (**Fig. 1d, e**), consistent with the idea that KIF18A inhibitor sensitivity is linked to the ability of tumor cells to sustain a checkpoint-dependent mitotic arrest.

### Rb pathway inactivation is associated with vulnerability to KIF18A inhibition

Because the Rb-E2F axis is a major regulator of mitotic gene expression, and MAD2 is a direct E2F target^15^, we reasoned that Rb-pathway state might distinguish the subset of ovarian cancer models capable of sustaining KIF18A inhibitor-induced mitotic arrest. We therefore profiled Rb-pathway proteins across ovarian cancer cell lines spanning high, intermediate and low sensitivity to VLS-1488 treatment. The most sensitive models showed a recurrent pattern characterized by reduced total Rb protein, reduced phospho-Rb, and increased p16INK4a (**Fig. 2a,b**). CDKN2A, which encodes p16INK4a is a direct target of E2F^19^, and represents a potential for negative feedback inhibition, as p16INK4a activates Rb through inhibition by the formation of Cyclin D/CDK4^20^. However, in cancer, this negative feedback loop is futile and as such increased p16INK4a expression is widely considered a marker for Rb pathway inactivation^21^.

**Fig. 2.**
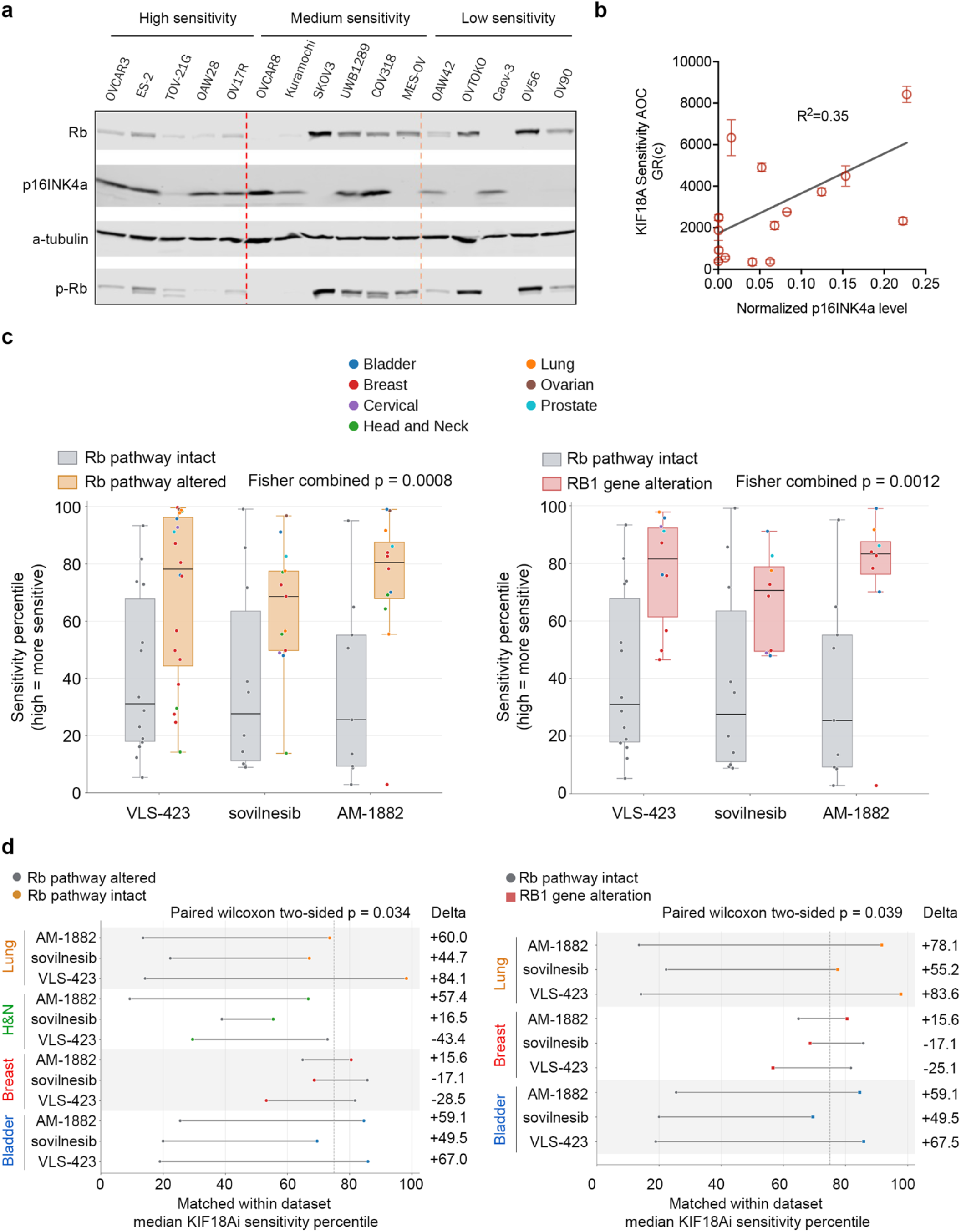
An Rb-low, p16INK4a-high ovarian cancer cell state is associated with KIF18A inhibitor sensitivity. **a**, Immunoblot analysis of Rb, p16INK4a and phospho-Rb across ovarian cancer cell lines grouped by KIF18A inhibitor sensitivity. α-tubulin served as a loading control. **b**, Correlation between normalized p16INK4a protein levels and KIF18A sensitivity score based on GR(c) across the ovarian cancer cell line panel. **c**, Rb pathway disruption is associated with increased KIF18A inhibitor sensitivity across independent pan-cancer datasets. *Left*, models with broad Rb pathway alteration, defined by direct RB1 defect, CCNE1 amplification or Rb hyperphosphorylation, or CDKN2A deletion or inactivating mutation, were compared with Rb pathway intact models across the VLS-423, sovilnesib, and AM-1882 datasets. Right, the altered group was restricted to models with direct RB1 gene alteration. Responses were harmonized to a common sensitivity-percentile scale such that higher values indicate greater sensitivity. Tukey box plots show the distribution of sensitivity percentiles within each dataset, and individual points represent cell lines colored by histology. Across all three datasets, both broad Rb pathway alteration and direct RB1 alteration were associated with a consistent shift toward greater KIF18A inhibitor sensitivity, with the strongest and most reproducible effect observed for direct RB1 alteration. P values were combined across the three independent datasets using Fisher’s method. **d**, Cancer type-matched analysis of KIF18A inhibitor sensitivity as a function of Rb pathway status. *Left*, models with broad Rb pathway alteration, defined by direct RB1 defect, CCNE1 amplification or Rb hyperphosphorylation, or CDKN2A deletion or inactivating mutations, were compared with matched Rb pathway intact models within the same histology and dataset. *Right*, the altered group was restricted to models with direct RB1 gene alteration. Each line connects the median sensitivity percentile of the intact and altered groups within an evaluable histology-dataset pair. Positive shifts indicate greater sensitivity of the altered group. Broad Rb pathway alteration was associated with a median gain of 47.1 percentile points across 12 evaluable matched comparisons (P = 0.034), whereas direct RB1 alteration was associated with a median gain of 55.2 percentile points across 9 matched comparisons (P = 0.039). The effect was especially consistent in bladder and lung.

To ask whether more generic downstream effects of chromosomal instability were similarly informative at separating KIF18A inhibitor-sensitive from insensitive cell lines, we quantified the basal levels of micronuclei across the same ovarian panel. Most of the cell lines exhibited elevated basal frequencies of micronuclei in line with ovarian cancer being a predominantly chromosomally unstable cancer type (**Supplementary Fig. 3a**). However, micronuclei burden did not correlate with drug sensitivity (**Supplementary Fig. 3b**). Similarly, there were no discernable differences in the frequencies of chromosome segregation defects – defined as lagging chromosomes, chromatin bridges, or multi-polar mitoses – between KIF18Ai sensitive and insensitive cell lines. Therefore, within a disease setting broadly enriched for chromosomal instability, KIF18A inhibitor sensitivity is associated not simply with CIN in general, but likely with a subset of chromosomally unstable cancers with Rb pathway defects.

We then tested whether p16INK4a expression could be captured using a clinically implementable immuno-histochemical assay. In a cell-pellet tissue microarray, p16INK4a immunohistochemical intensity tracked with sensitivity to VLS-1488 and correlated with ly-sate-based p16INK4a abundance by immunoblot (**Supplementary Fig. 4**), supporting p16INK4a as a practical surrogate for the underlying biology in a manner that could be clinically feasible.

We next asked whether the Rb-linked sensitivity state uncovered in the ovarian cancer cell lines reflects a broader vulnerability to KIF18A inhibition across other cancer types. To address this, we performed a retrospective analysis across three independent pan-cancer sensitivity datasets generated with distinct KIF18A inhibitors, VLS-423, sovilnesib, and AM-1882^13^, comprising 321, 893, and 631 cancer cell lines, respectively. None of these screens was designed around the Rb pathway, making them a stringent test of whether Rb status predicts response beyond lineage. Responses were harmonized to a common sensitivity-percentile scale, and Rb altered and intact models were first compared within each dataset irrespective of histology. Rb status was defined using explicit cell line-level evidence of pathway disruption, including direct RB1 gene loss or inactivating mutation, CCNE1 amplification or Rb hyperphosphorylation, or CDKN2A deletion, and was also analyzed separately using the more restrictive category of direct RB1 gene alteration alone.

Under these conditions, models with broad Rb pathway disruption showed a consistent shift toward greater KIF18A inhibitor sensitivity across the three datasets, with significance supported by Fisher’s method for combining P values (**Fig. 2c**, *left*). This relationship became even stronger when the altered group was restricted to models with direct RB1 gene loss or inactivating mutation, with the most pronounced and reproducible shift observed in this subset (**Fig. 2c**, *right*).

We next asked whether the link between Rb deficiency and KIF18A inhibitor sensitivity would persist after controlling for tumor lineage. We therefore carried out a histology-matched analysis in which altered and intact models were compared only within the same cancer type and within the same screen. Broad Rb pathway inactivation, encompassing direct RB1 alteration, CCNE1 amplification or Rb hyperphosphorylation, and CDKN2A deletion or mutation, remained associated with increased KIF18A inhibitor sensitivity, with Rb pathway alterations showing increased sensitivity to KIF18A inhibitors in 9 of 12 evaluable comparisons, each comprised of groups of Rb-proficient and Rb-deficient cell lines in the same cancer type, and showing a median increase of 47.1 percentile points in the ranked sensitivity score (**Fig. 2d**, *left*; P = 0.034). Similarly, limiting the analysis to direct RB1 gene alteration made the effect even more pronounced, with positive shifts in 7 of 9 cancer type-matched comparisons and a median increase of 55.2 percentile points (**Fig. 2d**, *right*; P = 0.039).

The recurrence of this pattern across multiple lineages, most consistently in bladder and lung, argues that the KIF18A-sensitive Rb-deficient state is a cross-histology biological phenotype rather than an ovarian-restricted or lineage-driven effect. More generally, the breadth and reproducibility of this effect is notable given that the three datasets were generated independently and with different KIF18A inhibitors sharing a similar mechanism of action yet converged on the same biological relationship.

### Active spindle assembly checkpoint signaling is required for KIF18Ai sensitivity in Rb-low cell lines

We next asked whether a functional spindle assembly checkpoint was required for response to KIF18A inhibition in Rb-deficient models. To test this, we disrupted Mad1 function using CRISPR-Cas9 in otherwise KIF18A inhibitor-sensitive and Rb-deficient OVCAR3 and OVCAR8 ovarian cancer cell lines. We also deleted MAD1 in BT549, a triple-negative breast cancer cell line known to have RB1 genomic loss^22^. In each background, MAD1 knockout was sufficient to confer resistance to VLS-1488 treatment (**Fig. 3a,b**), consistent with the hypothesis that deficiency in Rb function promotes sensitivity to KIF18A inhibitors in a manner that is dependent on a functional spindle assembly checkpoint. Taken together, these data identify an Rb-low, p16INK4a-high ovarian cancer cell state associated with KIF18A inhibitor sensitivity and support a model in which Rb-pathway disruption marks tumors capable of sustaining checkpoint-dependent mitotic arrest required for drug response.

**Fig. 3.**
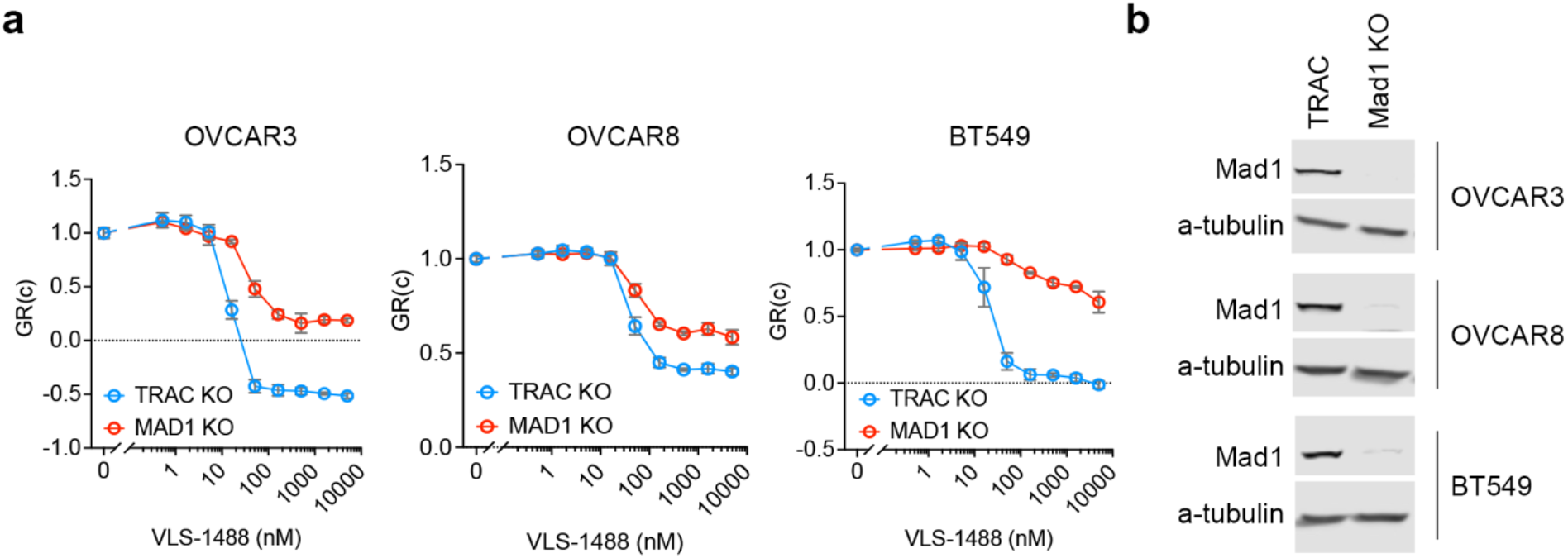
Disrupting spindle assembly checkpoint signaling in Rb-deficient cell lines promotes KIF18Ai resistance. **a**, Growth-rate dose-response curves for VLS-1488 in TRAC knockout control and their MAD1 knockout derivatives of OVCAR3, OVCAR8 and BT549 cells. **b**, Immunoblot confirmation of MAD1 knockout in the indicated cell lines. α-tubulin served as a loading control.

### p16INK4a as a clinically deployable assay to capture on-mechanism responder state across two KIF18A inhibitors

We next asked whether this Rb-linked responder state could be measured directly in patient pre-treatment tumor biopsies. To do so, we established centralized p16INK4a IHC on baseline tumor specimens from HGSOC cohorts treated on three clinical protocols: the ongoing phase I/II VLS-1488 study (NCT05902988), an ongoing dose-finding sovilnesib study (NCT06084416), and a completed dose escalation study of sovilnesib (NCT04293094). Because VLS-1488 and sovilnesib share the same mechanism of action^6,7^, this design allowed us to test whether the biomarker identified an on-mechanism responder state across two distinct KIF18A inhibitors rather than a compound-specific association. Tumors were classified as biomarker-positive by the presence of strong, diffuse 3+ staining in more than 90% of tumor cells (**Fig. 4a,b**), a threshold intended to identify the binary nature of the underlying mechanism of p16INK4a expression upon Rb loss.

**Fig. 4.**
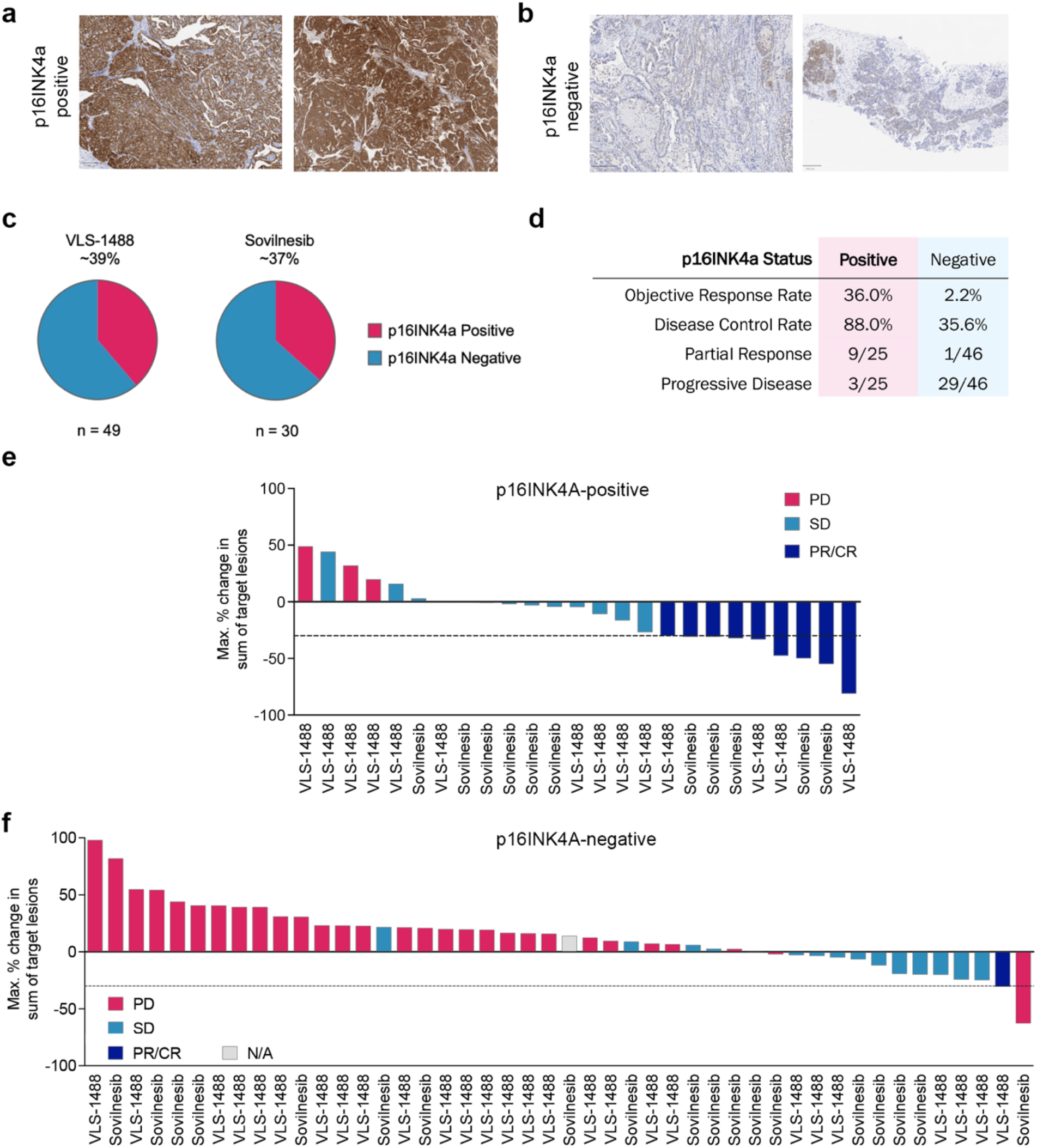
Strong p16INK4a staining identifies responders to KIF18A inhibition in high-grade serous ovarian cancer. **a**,**b**, Representative baseline pre-treatment tumor samples classified as p16INK4a-positive (a) or p16INK4a-negative (b) by IHC. p16INK4a positivity was defined as strong, diffuse 3+ staining in more than 90% of tumor cells. **c**, Prevalence of p16INK4a-positive tumors in the VLS-1488 and sovilnesib cohorts. Numbers below the pie charts indicate the number of analyzed samples in each cohort. **d**, Summary of ORR, DCR, partial response and progressive disease according to p16INK4a status in the pooled clinical dataset. **e**,**f**, Waterfall plots of maximum percentage change in the sum of target lesions in p16INK4a-positive (e) and p16INK4a-negative (f) tumors from patients treated with VLS-1488 or sovilnesib. Bar colors denote best overall response category. The dashed line indicates the RECIST threshold for partial response. One case with available post-baseline imaging is shown in grey was excluded from best overall response analyses for not meeting the pre-specified criteria for RE-CIST evaluability. In f, one patient met the RECIST threshold for target lesion shrinkage but was classified as having progressive disease at the first on-treatment assessment and was therefore scored as progressive disease for best overall response.

After pre-specified quality control, 79 samples were included in the biomarker prevalence analysis, 70 were evaluable for objective response, and 71 were evaluable for progression-free survival (**Fig. 4c, Table 1** and **Supplementary Table 2**). The analyzed set comprised 49 VLS-1488-treated samples and 30 sovilnesib-treated samples. p16INK4A positivity was observed in 19 of 49 analyzed VLS-1488 tumor samples and 11 of 30 analyzed sovilnesib tumor samples, corresponding to a pooled prevalence of 30 of 79 tumors, or ∼38%. Progression-free survival (PFS) data were not available for patients treated on NCT04293094, and sovilnesib PFS analyses were therefore restricted to cases with available follow-up.

**Table 1.** Response Summary Statistics.

| Cohort | Biomarker prevalence (%) | ORR evaluable Biomarker +/- (total) | Biomarker-positive ORR [95% CI] | Biomarker-negative ORR [95% CI] | Unselected ORR [95% CI] | Fisher exact p | Bi-omarker-Positive DCR | Bi-omarker-Negative DCR |
| --- | --- | --- | --- | --- | --- | --- | --- | --- |
| Combined (VLS-1488 and Sovilnesib) | 30/79 (38.0%) | 25/45 (total 70) | **9/25 (36.0%)**<br>[18.0% to 57.5%] | 1/45 (2.2%)<br>[0.1% to 11.8%] | 10/70 (14.3%)<br>[7.1% to 24.7%] | 0.0002 | 22/25 (88.0%) | 16/45 (35.6%) |
| VLS-1488 | 19/49 (38.8%) | 14/27 (total 41) | **4/14 (28.6%)**<br>[8.4% to 58.1%] | 1/27 (3.7%)<br>[0.1% to 19.0%] | 5/41 (12.2%)<br>[4.1% to 26.2%] | 0.0387 | 11/14 (78.6%) | 7/27 (25.9%) |
| VLS-1488 expansion-eligible ($\leq 5$ prior lines; 200–800 mg) | | 13/15 (total 28) | **4/13 (30.8%)**<br>[9.1% to 61.4%] | 0/15 (0.0%)<br>[0.0% to 21.8%] | 4/28 (14.3%)<br>[3.9% to 31.7%] | 0.0349 | 10/13 (76.9%) | 3/15 (20.0%) |
| Sovilnesib | 11/30 (36.7%) | 11/18 (total 29) | **5/11 (45.5%)**<br>[16.7% to 76.6%] | 0/18 (0.0%)<br>[0.0% to 18.5%] | 5/29 (17.2%)<br>[5.8% to 35.8%] | 0.0039 | 11/11 (100.0%) | 9/18 (50.0%) |

Caveats and assumptions: Biomarker definition: p16INK4a IHC strong and diffuse (3+) in >90% of tumor cells | Sovilnesib samples represent an aggregate of samples obtained from the Amgen sovilnesib/AMG650 trial (n = 8) in addition to ones from the Volastra Sovilnesib trial | ORR confidence intervals are Clopper-Pearson exact 95% binomial CIs | ORR group comparison p-values are two-sided Fisher exact tests on responder vs non-responder 2x2 tables | No adjustment for multiplicity, covariates (dose, prior lines, platinum status), or site effects is applied here. These are retrospective, hypothesis-generating analyses | Interpretation risk: biomarker appears predictive, but retrospective selection can inflate effect size; prospective validation is required | DCR: Disease Control Rate, ORR: Objective Response Rate

### Strong p16INK4a expression enriches for objective response across VLS-1488 and sovilnesib

To determine whether this pathology-defined state identified clinical responders, we first examined objective radiographic response in the pooled response-evaluable dataset. p16INK4a-positive tumors showed a markedly higher objective response rate (ORR) than p16INK4a-negative tumors, with responses in 9 of 25 (36%) biomarker-positive cases, as compared with 1 of 45 (2.2%) biomarker-negative cases (**Fig. 4d-f** and **Table 1**). Disease control rate (DCR) was likewise enriched in the p16INK4a-positive group, 22 of 25 cases, 88.0%, versus 16 of 45 cases, 35.6%. The distribution of maximal target-lesion change was shifted strongly toward tumor shrinkage in the p16INK4a-positive sub-group, with nearly all objective responses arising in biomarker-positive tumors (**Fig. 4e,f** and **Supplementary Fig. 5**).

This enrichment was reproduced across both drugs. In the sovilnesib cohort, responses were observed in 5 of 11 p16INK4a-positive tumors, 45.5%, whereas no responses were observed among 18 p16INK4a-negative tumors (**Fig. 5a** and **Table 1**). In the VLS-1488 cohort, objective responses were observed in 4 of 14 p16INK4a-positive tumors, or 28.6%, compared with 1 of 27 p16INK4a-negative tumors, 3.7% (**Fig. 5b** and **Table 1**). In a combined multivariable logistic regression model incorporating p16INK4a status, study drug and dose cutoff (high dose defined as 400 mg QD or greater for VLS-1488 and 100 mg QD or greater for sovilnesib), p16INK4a status remained the variable most strongly associated with objective response, whereas adjustment for study drug and dose did not account for the observed biomarker-response association (**Fig. 5c**). This effect was preserved in a subset analysis that additionally adjusted for number of prior lines of therapy (**Fig. 5d**). Notably, responses among p16INK4a-positive VLS-1488-treated tumors were observed across multiple active dose levels, arguing against a narrow drug exposure window as the explanation for the association (**Fig. 5b** and **Supplementary Fig. 6a**). This is further supported by the durable responses in p16INK4a-positive patients treated with 50 mg QD of sovilnesib and 200 mg QD of VLS-1488 (**Supplementary Fig. 6b-c**).

**Fig. 5.**
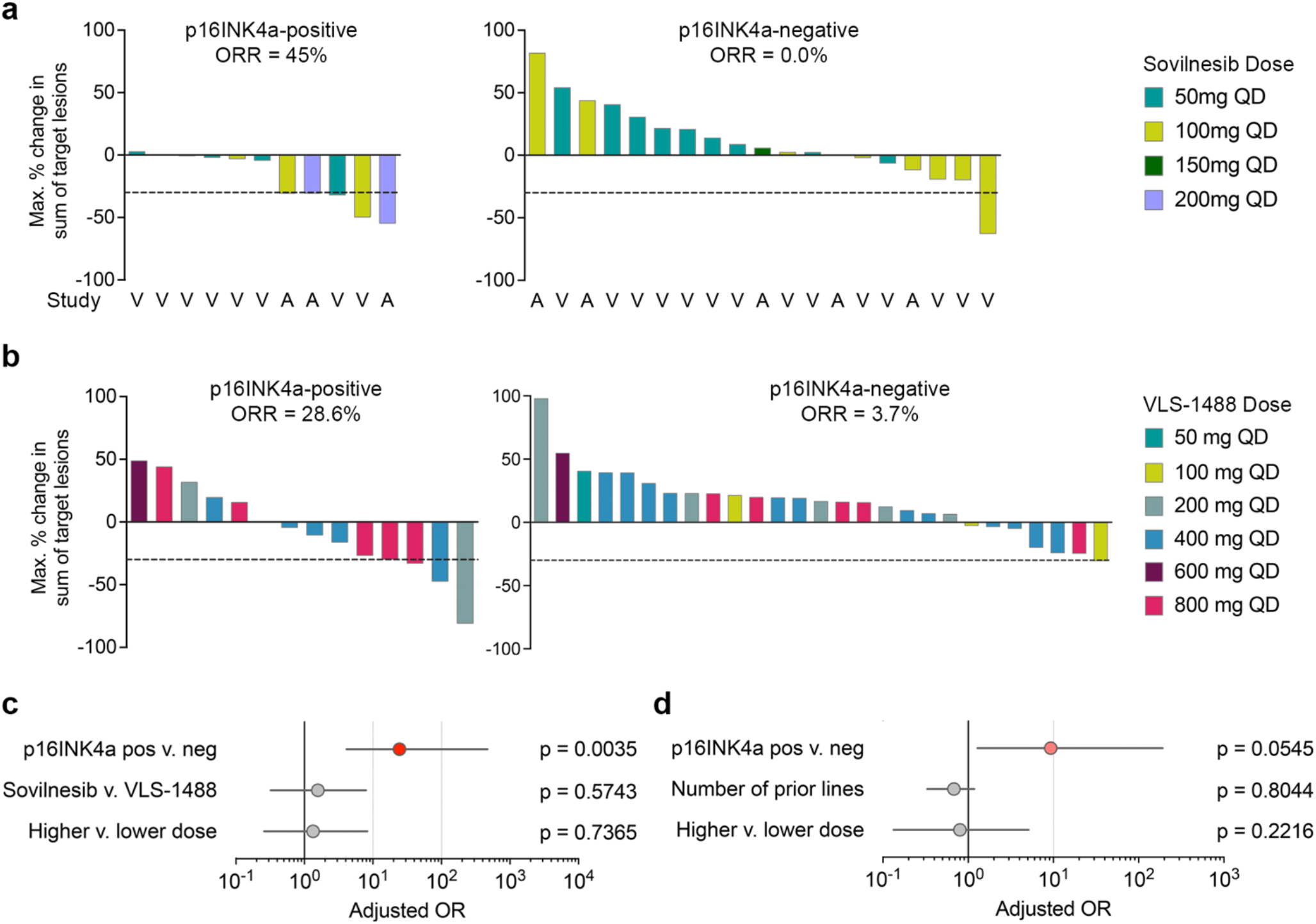
The association between p16INK4a and objective response is reproduced across VLS-1488 and sovilnesib. **a**, Waterfall plots of maximum percentage change in the sum of target lesions for sovilnesib-treated high-grade serous ovarian cancers, stratified by p16INK4a status and colored by dose level. Study origin is indicated on the x axis (A, Amgen-sponsored protocol, NCT 04293094; V, Volastra-sponsored protocol NCT06084416). **b**, Waterfall plots for VLs-1488-treated high-grade serous ovarian cancers, stratified by p16INK4a status and colored by dose level. **c**, Multivariable logistic regression analysis of objective response in the combined sovilnesib and VLS-1488 cohorts, adjusting for p16INK4a status, study drug and dose; higher dose defined as 400mg QD or above for VLS-1488 and 100 mg QD or above for sovilnesib. **d**, Sensitivity multivariable logistic regression analysis of objective response in the VLS-1488 cohort, adjusting for p16INK4a status, dose and number of prior lines. Odds ratios and 95% confidence intervals are shown.

### Strong p16INK4a expression is associated with durable benefit and prolonged progression-free survival

We next asked whether the response enrichment associated with p16INK4a translated into more durable clinical benefit. In the pooled cohort, p16INK4a-positive tumors were associated with substantially longer progression-free survival than p16INK4a-negative tumors, with median progression-free survival of 24.3 weeks versus 7.9 weeks and a hazard ratio for progression of 0.16 (**Fig. 6a** and **Table 2**). A similar pattern was observed in the sovilnesib subset with available PFS data, with medians of 39.7 weeks and 8.0 weeks in p16INK4a-positive and p16INK4a-negative tumors, respectively (**Fig. 6b** and **Table 2**). Likewise, this separation was reproduced in the VLS-1488 cohort, in which median progression-free survival of the p16INK4a-positive cohort was 24.0 weeks (**Fig. 6c** and **Table 2**). Importantly, PFS differences between p16INK4a-positive and p16INK4a-negative remained consistent even within the subset of patients treated with sovilnesib at the 50 mg QD assigned dose (**Fig. 6d**).

**Table 2.**
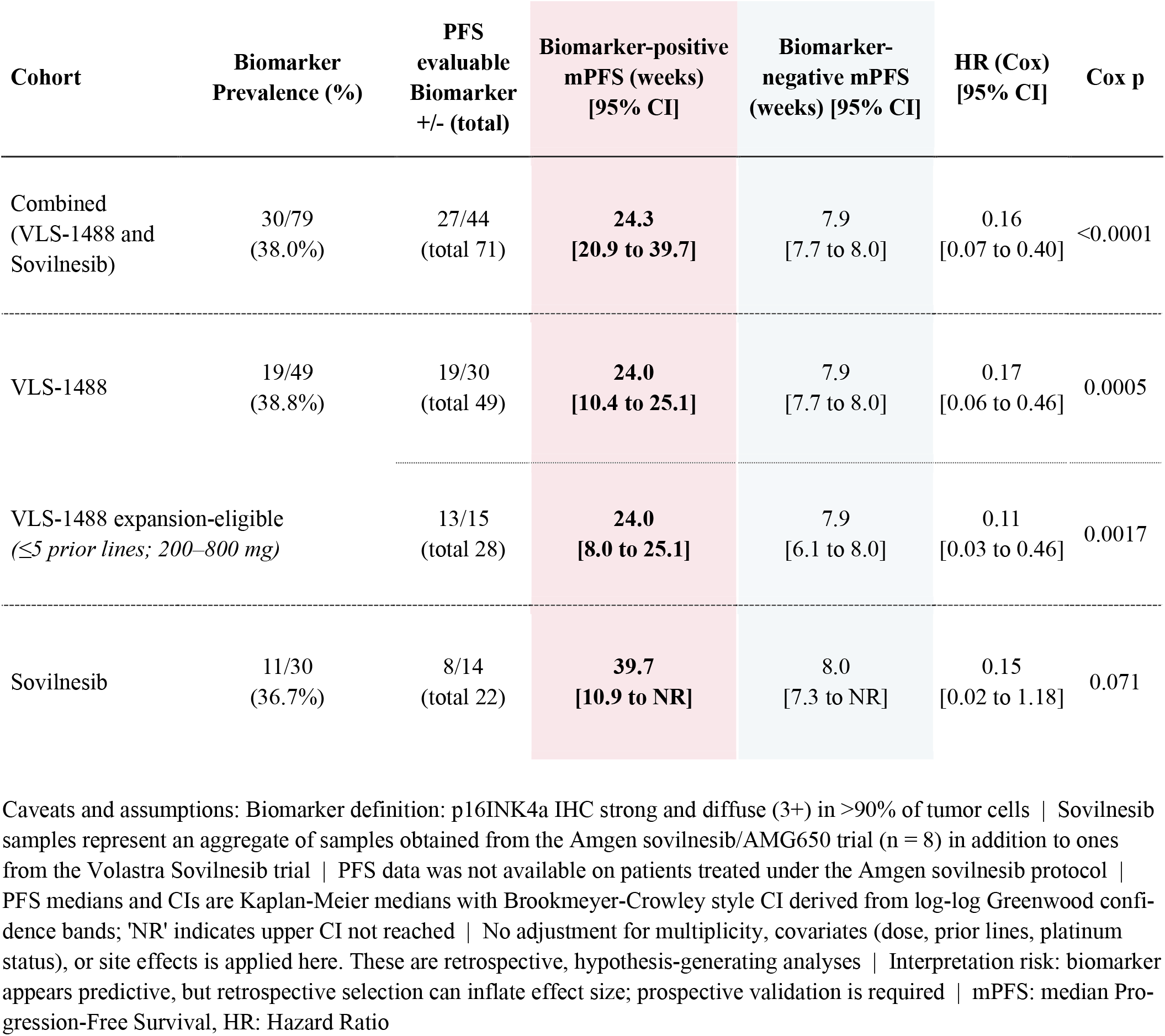
Progression-Free Survival Summary Statistics.

**Fig. 6.**
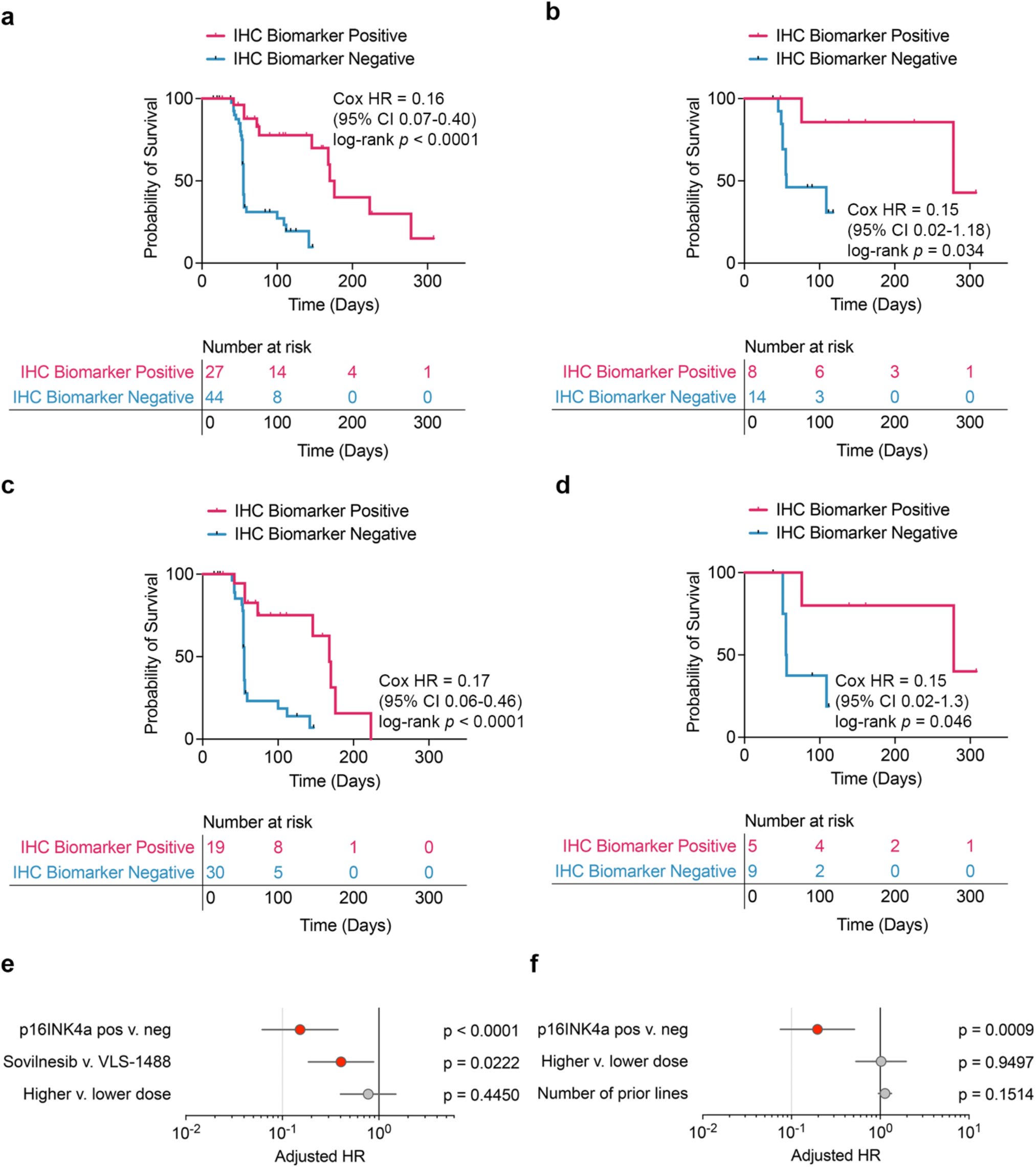
p16INK4a positivity is associated with prolonged progression-free survival upon KIF18A inhibition. **a-d**, Kaplan-Meier estimates of progression-free survival according to p16INK4a status in the combined cohort (a), the sovilnesib-treated cohort (b and d with d limited to patients treated with 50 mg QD of sovilnesib) and the VLS-1488-treated cohort (c) with available progression-free survival data. Tick marks denote censored observations. Numbers at risk are shown below each plot. **e**, Multivariable Cox regression analysis of progression-free survival in the combined cohort, adjusting for p16INK4a status, study drug and dose with high dose defined as 100 mg or above for sovilnesib and 400 mg and above for VLS-1488. **f**, Sensitivity multivariable Cox regression analysis of progression-free survival in the VLS-1488 cohort, adjusting for p16INK4a status, dose and number of prior lines. Hazard ratios and 95% confidence intervals are shown.

To test the robustness of this association, we performed multivariable Cox regression analyses. In both the combined dataset and the additional analysis incorporating prior lines of therapy, p16INK4a status remained independently associated with a lower risk of progression, whereas dose and the number of prior lines were not associated with progression-free survival in either model (**Fig. 6e,f**). Modeling p16INK4a as a continuous H-score yielded concordant results for both objective response and progression-free survival (**Supplementary Fig. 7**), indicating that the association was not dependent on a single binary cutoff. In the clinically relevant expansion-eligible VLS-1488 subset with 5 or fewer prior lines of therapy, and in which patients received doses ranging from 200 mg QD to 800 mg QD, strong p16INK4a expression also continued to enrich both radiographic activity and time on treatment (**Supplementary Fig. 6**), further supporting the robustness of the signal. Together, these findings define p16INK4a as a clinically deployable readout of an Rb-pathway-linked, spindle assembly checkpoint-competent responder state, and show that this state enriches for both response and durability across two KIF18A inhibitors with a shared mechanism of action.

## DISCUSSION

### p16INK4a identifies an on-mechanism responder state to KIF18A inhibition

This study addresses a central barrier to the clinical development of KIF18A inhibitors, namely the lack of a clinically meaningful predictive biomarker. By integrating mechanistic work in pre-clinical models with retrospective clinical data from the analysis of pre-treated tumor biopsies from three clinical studies enrolling HGSOC patients, we identify strong diffuse p16INK4a expression as a practical readout of an Rb-pathway-inactivated responder state and show that this state is associated with both ORR and prolonged PFS. Several aspects of the current study are notable. First, the biomarker is not merely correlative at the clinical level but is rather anchored in a biologic framework in which KIF18A inhibitor sensitivity depends on the ability of tumor cells to sustain a lethal mitotic arrest. Second, the clinical signal is reproduced across two distinct KIF18A small molecule inhibitors, VLS-1488 and sovilnesib, that share the same mechanism of action^6,7^. That concordance is important because it argues that the association is target-level and on-mechanism rather than compound-specific. More broadly, these findings move the field beyond the earlier observation that KIF18A dependence is enriched in chromosomally unstable, aneuploid, or whole-genome-doubled cancers, and begin to define the specific biologic subset within CIN that is most likely to respond^5–9^.

The mechanistic implications are equally important. A recurring theme in prior KIF18A work is that chromosomal or karyotypic abnormalities may be necessary but not sufficient for conferring sensitivity to KIF18A inhibition, and that successful KIF18A inhibitor response requires a mitotic state in which cells cannot readily escape spindle assembly checkpoint engagement. Our data support that framework and place the Rb pathway upstream of it. Prior work showed that Mad1 and Mad2 are a direct E2F target and are aberrantly expressed in cells with Rb pathway defects, providing a mechanistic basis for coupling RB1 loss to checkpoint competence and mitotic vulnerability^15^. More recent work has also shown that KIF18A inhibitor lethality depends on the balance between spindle assembly checkpoint signaling and mitotic exit, such that cells predisposed to persistent checkpoint signaling are uniquely sensitive^14^. Within that context, p16INK4a is a surrogate for Rb-pathway inactivation. That surrogate is biologically plausible because reciprocal Rb loss and p16 upregulation has long been recognized in human cancers, and elevated p16 expression is the result of a futile feedback loop in the setting of Rb-pathway disruption in cancer^16,17^.

A major strength of p16INK4a IHC detection as a biomarker for patient selection is that it is already embedded in routine pathology practice for in vitro diagnostic (IVD) use. p16INK4a IHC is widely used in the diagnosis of HPV-associated cervical squamous lesions and is also established in routine assessment of oropharyngeal squamous cell carcinoma, where it functions as a clinically relevant surrogate marker^23,24^. In contrast to genomic signatures that may ultimately prove informative but require more complex workflows, p16INK4a detection by IHC offers an immediately testable biomarker that can be deployed on archival tissue with standard immunohistochemical infrastructure. At the same time, the current study also highlights an important nuance: the therapeutic substrate is Rb-pathway inactivation, not p16 positivity per se. That distinction matters because Rb pathway disruption can arise through several distinct mechanisms, including RB1 genomic loss, cyclin E or cyclin D/CDK4 activation most often through genomic amplification, and viral oncogene-mediated Rb suppression.

### Rb-pathway inactivation broadens the therapeutic opportunity for KIF18A inhibition

These findings also point to a broader therapeutic potential that extends well beyond ovarian cancer. Tumors driven by Rb-pathway inactivation currently lack targeted therapies and remain a major unmet need in oncology. Therapeutic strategies that depend on an intact RB1 gene, most notably CDK4/6 inhibitors, lose activity once the pathway is disabled, and RB1 loss or p16-high states have both been associated with intrinsic or acquired resistance to CDK4/6 inhibition^25–27^. KIF18A inhibition therefore opens a conceptually distinct avenue: rather than requiring Rb function, it appears to exploit the mitotic liabilities created when the pathway is disrupted. This is particularly attractive because Rb pathway lesions are common across cancer, yet therapeutically underexploited. And, in addition to p16 status, genomic features such as RB1 loss or CCNE1 amplification can serve as orthogonal biomarkers for patient selection.

The ovarian cancer setting may nevertheless be especially important for early development of KIF18A inhibitors. In HGSOC, one biologically plausible route into the Rb-low, p16-high state is CCNE1 amplification, which is associated with primary treatment failure, poor survival, and relative independence from BRCA1 and BRCA2 deficiency^28,29^. More broadly, HR-proficient HGSOC remains a poor-prognosis subgroup with limited benefit from PARP inhibitors and an urgent need for new maintenance approaches^30^. This is where the tolerability profile of KIF18A inhibitors may become particularly consequential. Maintenance therapy in ovarian cancer places exceptional weight on long-term tolerability, convenience, and the ability to preserve quality of life, and those considerations are especially relevant for patients who are PARP inhibitor-ineligible or unlikely to benefit from PARP inhibition. A well-tolerated oral KIF18A inhibitor directed to an Rb-pathway-inactivated subset could therefore have value not only in platinum-resistant disease, where the current data were generated, but also earlier in the disease course, particularly in HR-proficient and biologically Rb-dysregulated tumors. Importantly, the argument for prediction is strengthened by the fact that high p16INK4a expression is not prognostic in ovarian HGSOC in large independent datasets, reducing the likelihood that the present findings simply identify a biologically favorable subgroup^31^.

At the same time, the data suggest two parallel clinical paths. One is biomarker-selected development in tumor types where strong diffuse p16 is present in a meaningful but non-universal proportion of cases, as in HGSOC. The other is histology-enriched or biomarker-agnostic development in cancers where Rb-pathway disruption is already highly prevalent. Small-cell lung cancer is an obvious example, as the vast majority of cases are RB1-deficient^16,17^. HPV-associated cervical and oropharyngeal cancers are another, because functional Rb inactivation is central to their biology and p16 assessment is already part of standard pathologic practice^23,24^. Uterine serous carcinoma also warrants attention, given the high prevalence of strong diffuse p16 expression reported in that disease^32^. Together, these observations support a development strategy that couples biomarker-selected cohorts to tumor-type-selected cohorts, depending on whether Rb-pathway disruption is intermediate in prevalence, nearly universal, or better captured genomically than immunohistochemically.

### Prospective development, companion diagnostics, and study limitations

Several limitations should be emphasized. The clinical biomarker analyses were retrospective and were conducted across dose escalation and dose expansion cohorts, with inherent heterogeneity in dose, prior therapy, specimen availability, and evaluability. Accordingly, these data should not be used to directly compare the efficacy of VLS-1488 and sovilnesib, but rather to assess whether the biomarker association is directionally concordant across two inhibitors with a shared mechanism of action. Although the biomarker signal is consistent across both compounds and remains evident in multivariable models, the absolute effect size will require prospective confirmation. Moreover, because these cohorts were evaluated in an early phase setting, before dose optimization, and in heavily pretreated patients, the observed response estimates may underrepresent the activity achievable in prospectively selected populations treated at optimized dose and potentially earlier lines of therapy. The mechanistic data are also supportive rather than exhaustive. Our findings strongly argue that Rb pathway disruption and spindle assembly checkpoint competence are linked to KIF18A inhibitor response, but they do not yet prove that every p16 high tumor is sensitive or that every Rb disrupted tumor must be p16-high, an area of active investigation. That point is particularly relevant for pan cancer development, where the relationship among p16 expression, Rb function, viral oncogene status, and proliferative stress will vary by histology. The central translational challenge, therefore, is not simply to prospectively validate p16, but to define where p16 alone is sufficient, where it should be supplemented by RB1 or CCNE1 genomics, and finally where tumor histology itself may serve as the most efficient enrichment strategy.

A rational next step is a tiered prospective basket design. One tier could enroll p16-high tumors irrespective of histology using a pre-specified centralized IHC assay with the current cutoff. A second tier could focus on tumor-specific cohorts in which p16 prevalence and unmet need are both favorable, such as HGSOC and uterine serous carcinoma. A third tier could evaluate genomically or histologically selected cohorts with near-universal or alternative forms of Rb-pathway disruption, such as RB1-deficient small-cell lung cancer, CCNE1-amplified gynecologic cancers, or post-CDK4/6 inhibitor treated breast cancers with acquired Rb-pathway escape. This would establish p16INK4a as a rare kind of biomarker in mitotic therapy: mechanistically grounded, operationally tractable, and directly actionable at the point of care.

## MATERIALS AND METHODS

### Cell culture

HCC1806, OE19, OE33, TE1, TE8, TE9, TE10, TE15, KYSE30, KYSE70, KYSE140, KYSE180, KYSE410, KYSE450, KYSE520, and JHE-SOAD1 were cultured in RPMI (Gibco) supplemented with 10% FBS with 50 IU/mL penicillin and 50 ug/mL streptomycin. HET1A was cultured in BEGM (Lonza) supplemented with BEGM SingleQuot supplements and growth factors. CPA was cultured in KSFM (Gibco) supplemented with 5 g/L human recombinant epidermal growth factor and 50 mg/mL bovine pituitary extract. All cell lines were maintained at 37°C in a 5% CO_2_ atmosphere and routinely tested for mycoplasma contamination.

### Live-Cell Imaging of mitotic duration

To produce stably expressing H2B-GFP cell lines for live-cell imaging, cells were infected with H2B-GFP lentivirus generated from an H2B-GFP ORF cloned into the pLenti6/V5 lentiviral vector backbone (Invitrogen). H2B-GFP infected cell lines were then FACS sorted by GFP fluorescence. Stably expressing H2B-GFP cells were seeded on to glass-bottom 12-well tissue culture plates (Cellvis) and treated with either DMSO or KIF18Ai at the start of live-cell imaging (20,000 – 40,000 cells/well). Imaging was performed on a Nikon Eclipse Ti-E fluorescence microscope equipped with Nikon Perfect Focus Technology. The microscope was enclosed within a temperature and atmosphere-controlled environment maintaining 37°C and 5% humidified CO_2_. Brightfield and fluorescent images were captured every 5 minutes for 72 hours at multiple points per well using a 20X 0.45 NA Plan Fluor objective. Movies obtained from live-cell imaging were used to track individual cells through their respective cell cycles and record their mitotic durations and resulting fate (normal mitosis, micronuclei formation post-mitosis, death during mitosis, death post-mitotic slippage, and survival post-mitotic slippage). Mitotic duration was calculated as the time from the onset of chromosome condensation to either complete chromosome decondensation in daughter cells (for cells that completed mitosis) or to cell death (for cells that died during mitosis). Mitotic cells were analyzed within the first 24 hours to capture the first mitosis subjected to KIF18Ai and were tracked through to their next mitosis to determine cell fate. Mitotic duration and cell fate were recorded for ∼90 cells per condition per cell line across three biological replicates.

### Growth Inhibition Experiments

Growth inhibition was measured to determine the degree of KIF18Ai sensitivity across cell lines. Cells were seeded in triplicate in 96-well white-bottom plates (750-1000 cells/well). The following day, cells were treated with DMSO or KIF18Ai and cultured for an additional 6 days. To assess % growth inhibition, CellTiter-Glo 2.0 (Promega) was performed according to the manufacturer’s instructions and luminescence was measured with a BMG Clariostar plate reader. % growth inhibition was determined across three biological triplicates per cell line and was calculated using the following equation (LUM = luminescence signal obtained via CellTiter-Glo):

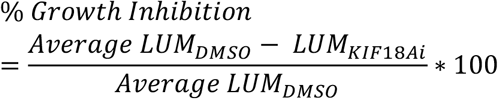

### Immunoblot of p16INK4a, Rb, phosphor RB, tubulin, MAD1

Cells were collected on Day 3 by centrifugation at 1500 rpm and frozen at -80C. For protein extract preparation, cell pellets were thawed on ice for 10 min and lysed by adding 50-100 ul of cell lysis buffer (50 uL 100X Protease/Phosphatase Inhibitor and 12.5 uL Benzonase (Sigma, R0278) at a 1:400 dilution to 5 mL of RIPA buffer). Resuspended cell pellets were placed on ice and incubated with lysis buffer for 10-15 min and vortexing during incubation. Lysates were cleared by spinning down at the 12700 rpm for 12 min at 4 C. After centrifugation, cleared cell lysates were transferred into new 1.5mL labeled tubes. Protein concentration was measured for each cell lysate, following the manufacturer’s protocol of Precision Red Advanced Protein Assay (Cytosketon, ADV02). Briefly, 3 uL of each cell lysate was added into each well of a clear 96-well culture plate, followed by 300 uL of Precision Red reagent. The plate was incubated on a shaker for 1 min, and luminance was measured with the Nivo Multimode Microplate Reader (Revvity) using established program. Cell lysates were used to prepare equal amount of protein (40 ug) for gel loading by adding proper amount of 4x LDS sample buffer, 10x reducing agent, Mili-Q water and cell lysate in a new correspondingly labelled 1.5 mL tube. Samples were denatured at 85C for 10 min on a hot plate and loaded on a 4-20% gel (Novex 4-20% Bis-Tris Gel, Invitrogen, WXP42020BOX). Gel was run using 80V for 20-30 min and then switched to 120V until the ladders are visually well separated. Gels were transferred to nitrocellulose membranes and blocked with LI COR Intercept in Tris Buffer Saline blocking buffer (TBS) (LI-COR, 92760001) at room temperature for 1 hr. Primary antibodies were prepared using LI COR Intercept TBS Blocking Buffer buffer and 0.2% Tween-20 (TBST). Membranes were incubated overnight with primary antibodies in blocking buffer at 4C on a rocker. The following antibodies and dilutions were used: total Rb antibody (Cell Signaling Technology, 9309S, 1:2000), CDKN2A antibody (Abcam, ab270058, 1:1000), anti a-tubulin (DM1A) (Cell Signaling Technology, 3873S, 1:1000), Phospho-Rb antibody (Cell Signaling Technology, 8516S, 1:1000), MAD2 antibody (Thermo Fisher, MA15832, 1:1000), MAD1(ThermoFisher PA5-28185, 1:1000), MAD1(Abcam, EPR14676-23, 1:1000). After the incubation, membranes were washed on a rocker with 1x TBST (BioWorld, 40120065) three times at RT for 5 min each time. Secondary antibodies were prepared as follows: goat anti-rabbit IR Dye 700CW (LI-COR, 926-32211) and goat anti-mouse IR Dye 800CW (LI-COR, 926-32210) in LI-COR TBS blocking buffer with 0.2% Tween-20. Membranes were incubated at RT for 1 hr on a rocker. Membranes were washed on a rocker with 1x TBST for four times at RT for 5 min each time. All membranes are imaged on the Odyssey CLx imager (LI-COR, Software version CLS-2693) machine.

### In vitro treatment of cell lines with VLS-1488 and GRc calculation

On the previous day of treatment, cells were harvested by trypsinization cells with 0.25% Trypsin-EDTA for 3 min at 37C, then neutralized and washed with full medium. Cells were spun at 1500 rpm for 3 min, resuspended in 5 mL of full medium and counted before plating. Four CTG plates were set up for each cell line for the four days of assessment (Day 0, 2, 4 and 6). One thousand 1 cells were seeded per well in 100 uL of media. On the day of the treatment, a master drug plate containing 21x drug serial dilutions was prepared. Briefly, a muti-channel pipette was used to add 54.3 uL DMSO into columns 4-11 in row C, E of a V bottom plate. Eighty ul of 10 mM VLS-1488 stock (diluted in DMSO) was added to C3 and 25.3 uL of the VLS-1488 stock to C4 and mixed well. After mixing, 25.3 uL was then transfered to next column 5, serially diluted to column 6; and this was repeated until column 11. The master drug plate was then used to dilute in media for a final dilution of 1:95. This was achieved by adding 188 ul of medium to column 2-11 for row C-F in a new V bottom plate and with the use of a multi-channel pipette, 2 uL of VLS-1488 dilutions from row C of the master drug plate were added to row C and mixed well. Two uL of DMSO was transferred from row G into column 11 from the 21x dilution plate. Finally, 5ul of drug was added from the intermediate plate to the cells using an automated pipette. Plates were placed on shaker to shake for 20 min. CellTiter Glo Luminescent Cell Viability Assay (Promega, G7570) was used on Day 0, 2, 4 and 6 following manufacturer’s instructions. GR(c) was calculated as described in Elizabeth et al.^33^ GR (c) = 2 (log_2_CTG_treated_/CTG_t0_)/ (log_2_CTG_untreated_/CTG_t0_)-1

### Pan-cancer KIF18A inhibitor response datasets

Three independent cancer cell-line response datasets were analyzed. The first was an internally generated VLS-423 response matrix, in which higher VLS-423 response values indicate greater sensitivity. The second was an sovilnesib pooled cell-line viability screen on 893 pan-cancer cell lines. The third was a AM-1882 dataset reported by Payton et al., which used a 5-day growth assay in DNA-barcoded pooled cell lines and determined response values for 631 cancer cell lines. Cell-line names were harmonized across datasets by upper-case normalization and alias collapsing to canonical CCLE or DepMap identifiers. Histology labels were assigned from canonical disease annotations. A curated Rb pathway status atlas was then assembled at the unique cell-line level using only explicit line-level evidence. Rb pathway intact status required affirmative evidence of intact Rb biology and was assigned from published functional Rb-positive lines in Mizuarai et al.^34^ or intact-like pRb expression/phosphorylation patterns in Robinson et al.^35^. Broad Rb patway-altered status required explicit evidence for one of three mechanisms: direct RB1 defect, defined as homozygous RB1 deletion, nonsense, frameshift or in-frame deletion, exon-deleting mutation, or published loss of RB protein/function; CCNE1 amplification / pRb-hyperphosphorylated Rb-off state; or CDKN2A / p16 lesion consistent with RB pathway disruption. These calls were curated from Mizuarai et al.^34^, Robinson et al.^35^, Scheffner et al.^36^, Baek et al.^37^, and Cellosaurus (https://www.cellosaurus.org/). Two discordant models, Hs578T and UM-UC-3, were excluded from binary comparisons because the database and literature annotations disagreed.

### Sensitivity harmonization and statistical analysis

Because the raw screening endpoints differed in direction across datasets, responses were harmonized to a common within-dataset sensitivity percentile scale, with higher percentiles always representing greater sensitivity. For VLS-423, percentiles were calculated directly from the raw response score, because higher raw values denote greater sensitivity. For sovilnesib and AM-1882, raw AUC values were reverse-ranked so that lower AUC corresponded to higher sensitivity percentile. Unmatched comparisons of Rb-defined groups within each dataset used two-sided Mann-Whitney U tests. Evidence across screens was summarized using Fisher’s method to combine P values. For histology-matched analyses, only dataset-histology strata containing both Rb-altered and Rb-intact models were retained. Within each matched stratum, effect size was defined as the difference between the median sensitivity percentile of the Rb-altered group and that of the Rb-intact group. These within-stratum deltas were then tested across strata with paired two-sided Wilcoxon signed-rank tests. For the direct RB1 analysis, the altered group was restricted to the direct RB1-defective subset while keeping the identical Rb-intact comparator. All tests were two-sided, and no unannotated cell line was treated as Rb-intact by default. Secondary descriptive analyses of responder enrichment used within-dataset quartiles, defined as the upper quartile of the VLS-423 score distribution and the lower quartile of the AUC distributions for sovilnesib and AM-1882. Analyses were performed in Python using pandas and SciPy.

### TRAC and MAD1 knock out cell lines generation

On Day 0 cells were harvested by trypsinization with 1.5 mL 0.25% Trypsin EDTA for 3 min at RT, neutralized and cells were washed with 4 mL of full medium. Cells were spun down at 1500 rpm for 3 min, resuspended with 4 mL medium, counted and diluted to concentration of 1×10^5^ cell/mL. Cells were plated a concentration of 2×10^5^ in each well of a 6 well (2 wells for each cell line). Cells were incubated cells at 37C for overnight. Two rounds of transfections were performed on Day 1 and Day 2, using 3 sgRNAs for each of the genes MAD1 sgRNA (Synthego, GCGUG-AGGUCGACCGCAACC, GAUAUUUCUAC-CUCGGCCCC, GCGUGAGGUCGACCGCAACC) and positive human sgTRAC (Synthego) using CRISPR max (Thermo Scientific, CMAX00008) and following manufacturer’s instructions. On Day 2, media was replaced in the morning to full media and in afternoon the transfection was repeated. On Day 3, transfection media was replaced to full media. Knock-down efficiency was tested by immunoblot after Day 4.

### Growth-rate dose-response curves for VLS-1488 in TRAC knockout control and their MAD1 knockout derivatives of OVCAR3, OVCAR8 and BT549 cells

On Day 0, OVCAR3, OVCAR8, and BT549 MAD1 KO cells and TRAC KO control were harvested by trypsinization with 0.25% Trypsin-EDTA for 3 min at 37C, neutralized and washed with full medium. Cells were spun down with 1500 rpm for 3 min and resuspended with 5 mL full medium and counted. Two thousand cells were plated per well in 100 uL of media and treatment was performed on Day 0 and 6 as described in section “In vitro treatment of cell lines with VLS-1488 and GRc calculation”

### Clinical Protocols and IRB approvals

**AMG-650 NCT04293094** is a phase 1, multicenter, open-label, dose-exploration and dose expansion study evaluating the safety, tolerability, pharmacokinetics, and efficacy of AMG650 (sovilnesib) in subjects with advanced solid tumors. 65 subjects dosed at doses ranging from 100mg QD to 300mg QD, including 29 subjects with HGSOC. **SOVI-2302 NCT06084416** is a Phase 1b dose optimization study of sovilnesib (an oral KIF18A inhibitor) in subjects with advanced high grade serous ovarian cancer. 47 subjects with HGSOC were dosed at 25mg (n=13) QD, 50mg QD (n=20) or 100mg QD (n=14). **VLS-1488-2201 NCT05902988** is a phase I/II Study of VLS-1488 (an oral KIF18A inhibitor) in subjects with advanced cancer. As of 5^th^ February 2026, had enrolled 140 subjects at dosed ranging from 50mg QD to 800mg QD, including 81 subjects with HGSOC. All clinical protocols, amendments, and informed consent forms for AMG-650, SOVI-2302, and VLS-1488-2201 were approved by the central IRB and by the institutional IRB or ethics committee at each participating site, as applicable, before study initiation. All studies were conducted in accordance with the Declaration of Helsinki, International Council for Harmonisation Good Clinical Practice guidelines, and applicable regulatory requirements.

### p16INK4a immunohistochemistry staining

Staining was performed using the Leica Bond RX automated stainer (Leica Microsystems) using a Standard Operating Procedure and fully automated workflow. Unstained slides were dewaxed using xylene and alcohol based dewaxing solutions. Epitope retrieval was performed by heat-induced epitope retrieval (HIER) of the formalin-fixed, using Tris based pH 9 solution (Leica Microsystems, AR9640) for 20 mins at 95 C. Tissues were first incubated with peroxide block buffer (Leica Microsystems), followed by incubation with the rabbit anti-CDKN2A (p16INK4a) antibody (Abcam, ab270058) at 1:1000 for 30 mins, followed by DAB rabbit secondary reagents: polymer, DAB refine and hematoxylin (Bond Polymer Refine Detection Kit, Leica Microsystems) according to the manufacturer’s protocol. The slides were dried, coverslipped (TissueTek-Prisma Coverslipper) and visualized using a Leica Aperio AT2 slide scanner (Leica Microsystems) at 40X.

### Statistical Analysis

Categorical variables were summarized as frequencies and percentages with 95% confidence intervals calculated using Wilson Score method. Association between categorical variables were assessed using Fisher’s exact test. Multivariate Logistic regression was performed to evaluate independent predictors of response outcome. Results are presented as adjusted odds ratios with 95% CI. The Kaplan-Meier method was used for survival estimation and the log-rank test was used for comparisons. Cox proportional hazards model was used for association analysis with survival outcome for univariate and multivariate analysis and for calculating the HR estimates along with 95% CIs. Two-sided P values of less than 0.05 were considered statistically significant. All statistical analysis was per-formed in R version 4.4.1 (2024-06-14).

## CONFLICTS OF INTEREST

C.A., A.A., S.C., A.R., S.B., N.B., D.P.S., T.B. and S.F.B. are employees of Volastra Therapeutics, from which they receive compensation and in which they hold equity. D.P.S. and S.F.B. serve as directors of Volastra Therapeutics. S.F.B., T.B., D.P.S., S.C. and S.B. are inventors on patent applications related to p16, RB-pathway inactivation, and KIF18A sensitivity in cancer.

## SUPPLEMENTARY FIGURES

**Supplementary Fig. 1.**
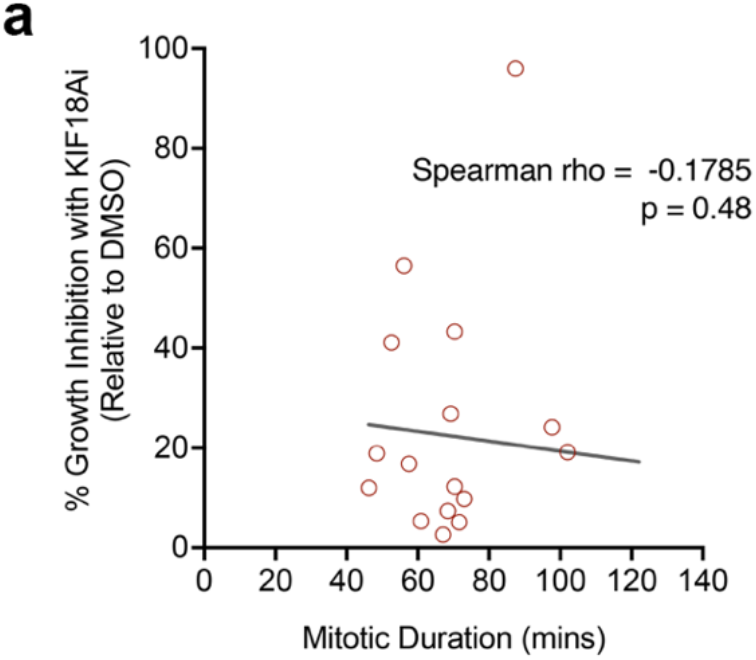
Baseline mitotic duration does not predict KIF18A inhibitor sensitivity. **a**, Correlation between baseline mitotic duration as measured by from live-cell imaging and growth inhibition following KIF18A inhibitor treatment in the esophageal cancer cell line panel.

**Supplementary Fig. 2.**
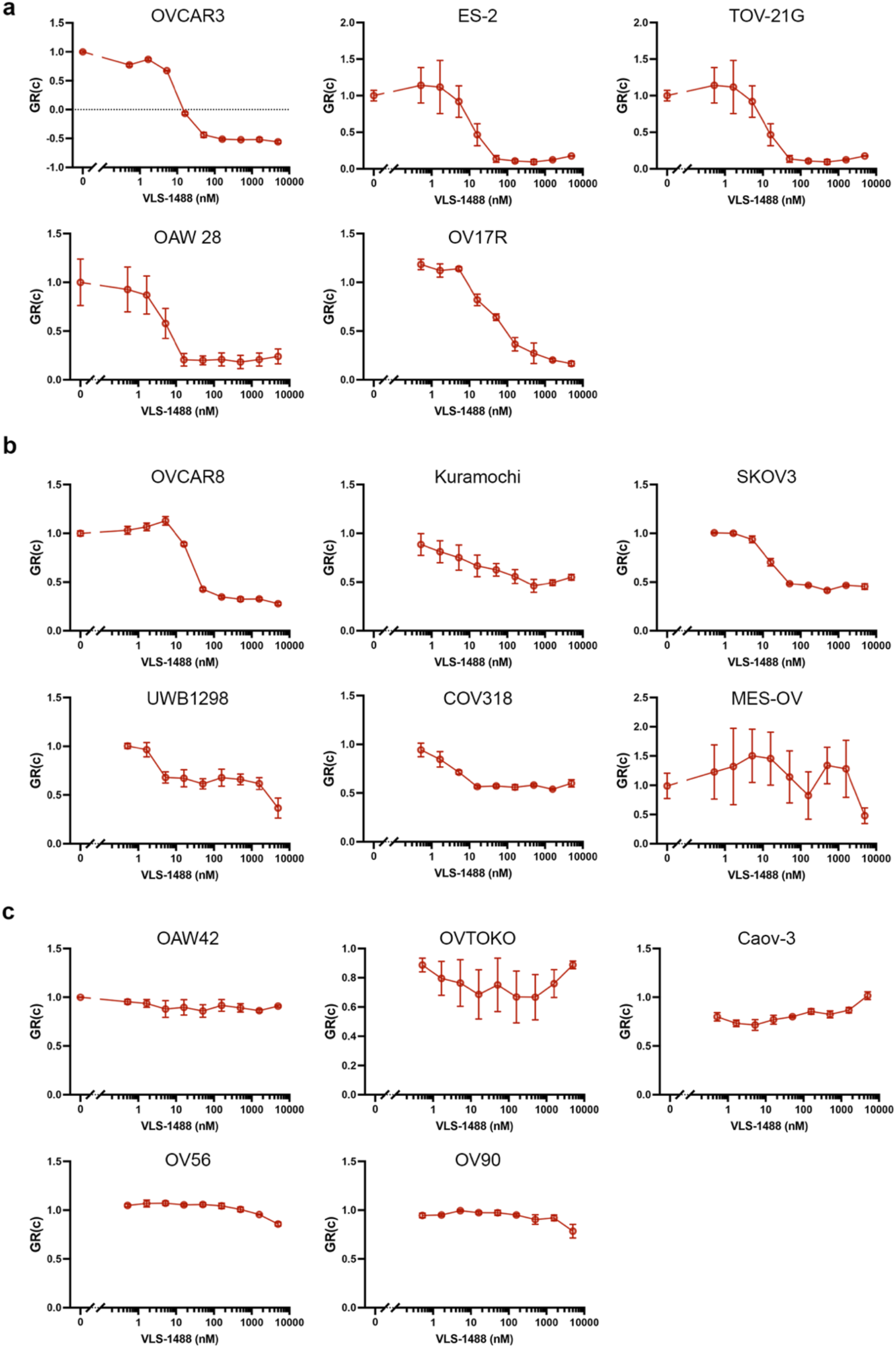
Ovarian cancer cell lines exhibit a range of sensitivities to VLS-1488, a KIF18A inhibitor. Growth rate inhibition curves, plotted as GR(c), for a panel of ovarian cancer cell lines treated with increasing concentrations of VLS-1488. Cell lines are grouped according to overall drug sensitivity. **a**, High-sensitivity ovarian cancer cell lines. **b**, Medium-sensitivity ovarian cancer cell lines. **c**, Low-sensitivity ovarian cancer cell lines. Data points represent mean values and error bars indicate s.d.

**Supplementary Fig. 3.**
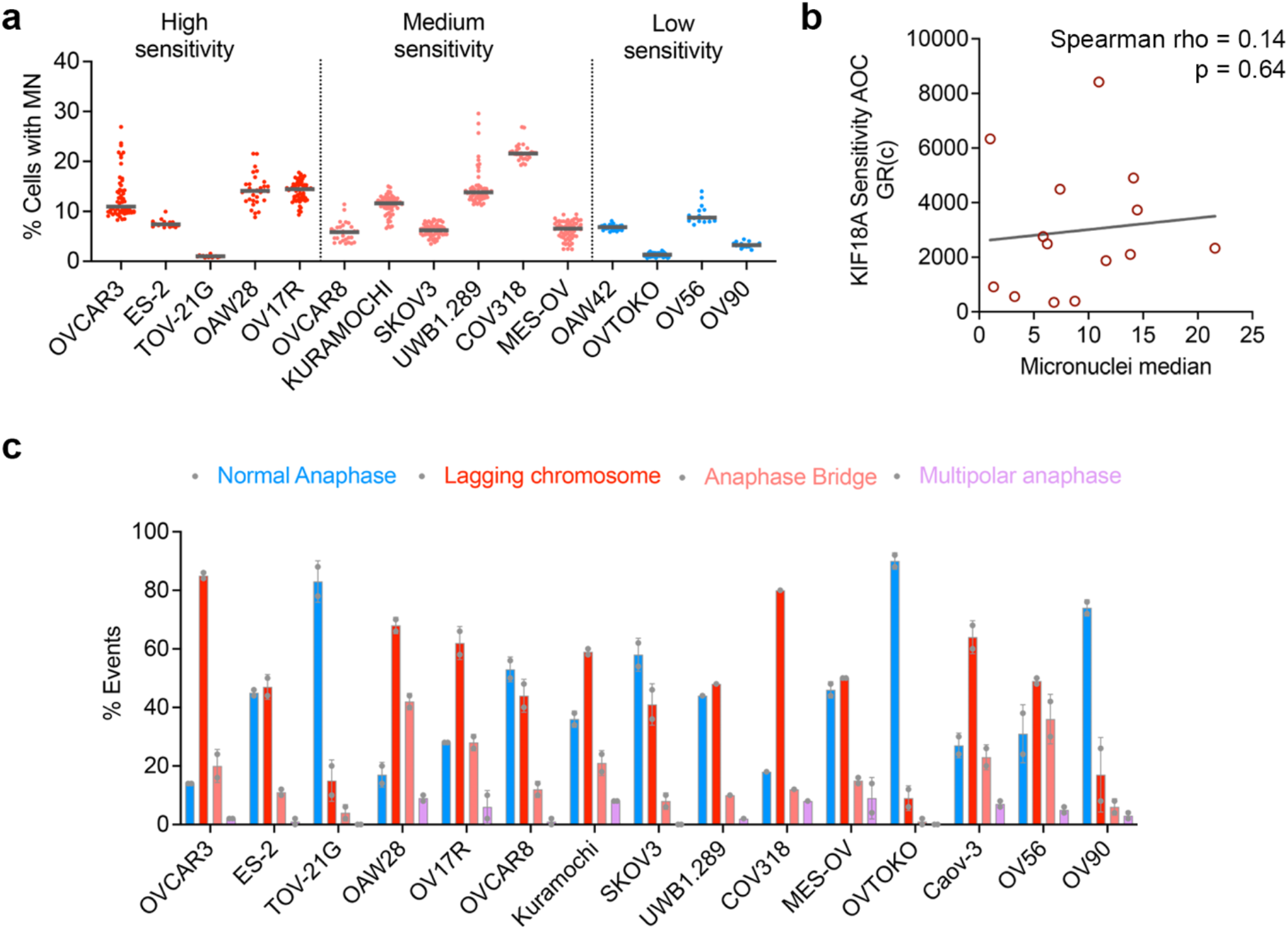
Micronuclei burden does not correlate with KIF18A inhibitor sensitivity in ovarian cancer cell lines. **a**, Frequency of cells with micronuclei across ovarian cancer cell lines grouped by KIF18A inhibitor sensitivity. Horizontal bars indicate the median. **b**, Correlation between median micronuclei frequency and KIF18A sensitivity across ovarian cancer cell lines. **c**, Frequency of cells with normal anaphase and abnormal anaphase (defined as anaphase with lagging chromosomes, chromatin bridges, or multipolar) across ovarian cancer cell lines grouped by KIF18A inhibitor sensitivity. Bars represent mean +/-SD.

**Supplementary Fig. 4.**
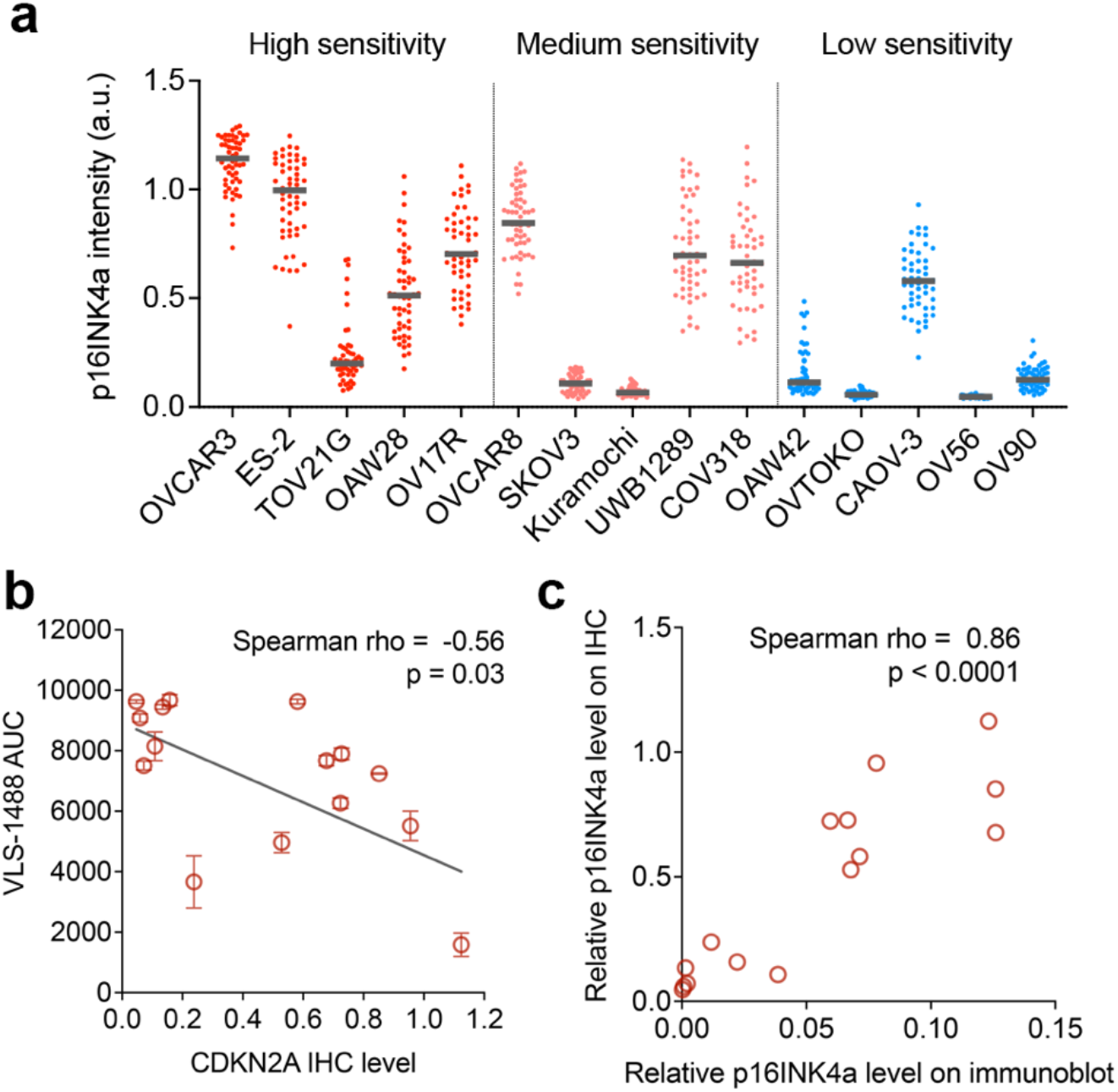
p16INK4a immunohistochemical intensity correlates with KIF18A inhibitor sensitivity and its immunoblot abundance in cell lysates. **a**, Quantification of p16INK4a immunohistochemical intensity in ovarian cancer cell-line tissue microarrays, grouped by KIF18A inhibitor sensitivity. Horizontal bars indicate the median. **b**, Correlation between CDKN2A IHC level and VLS-1488 sensitivity score. **c**, Correlation between p16INK4a abundance measured by IHC and by immunoblot.

**Supplementary Fig. 5.**
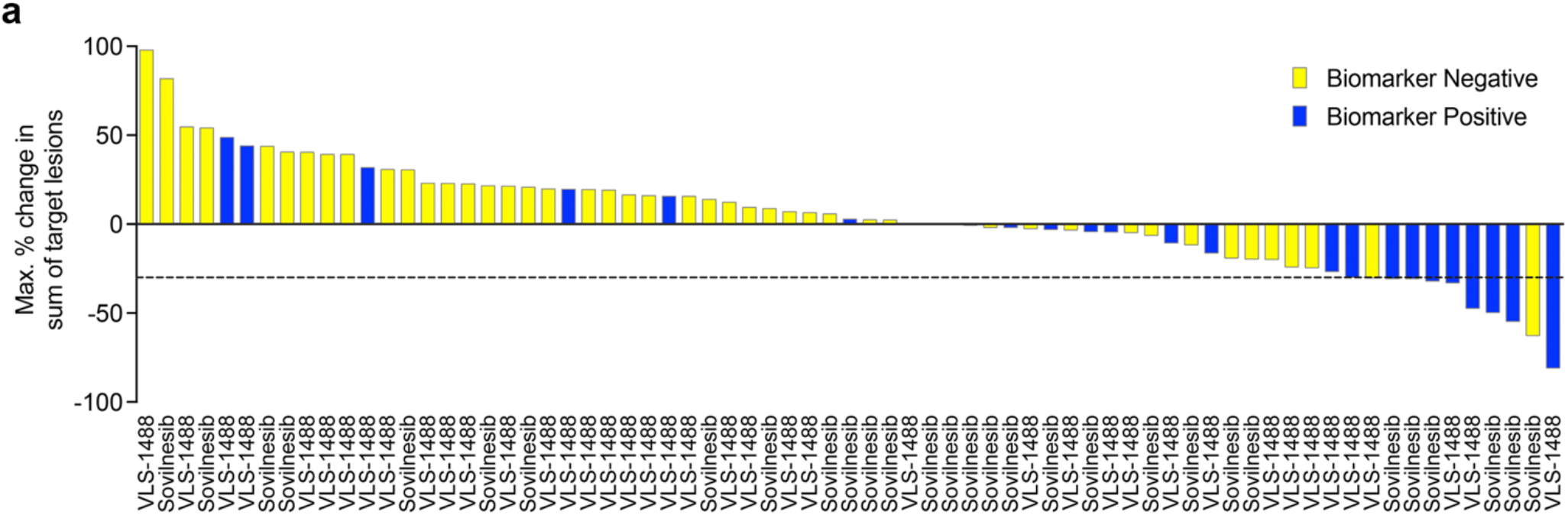
Distribution of p16INK4a biomarker status across the pooled response waterfall plot. **a**, Pooled waterfall plot of maximum percentage change in the sum of target lesions across the VLS-1488 and sovilnesib cohorts, with bars colored by p16INK4a biomarker status. The dashed line indicates the RECIST threshold for partial response.

**Supplementary Fig. 6.**
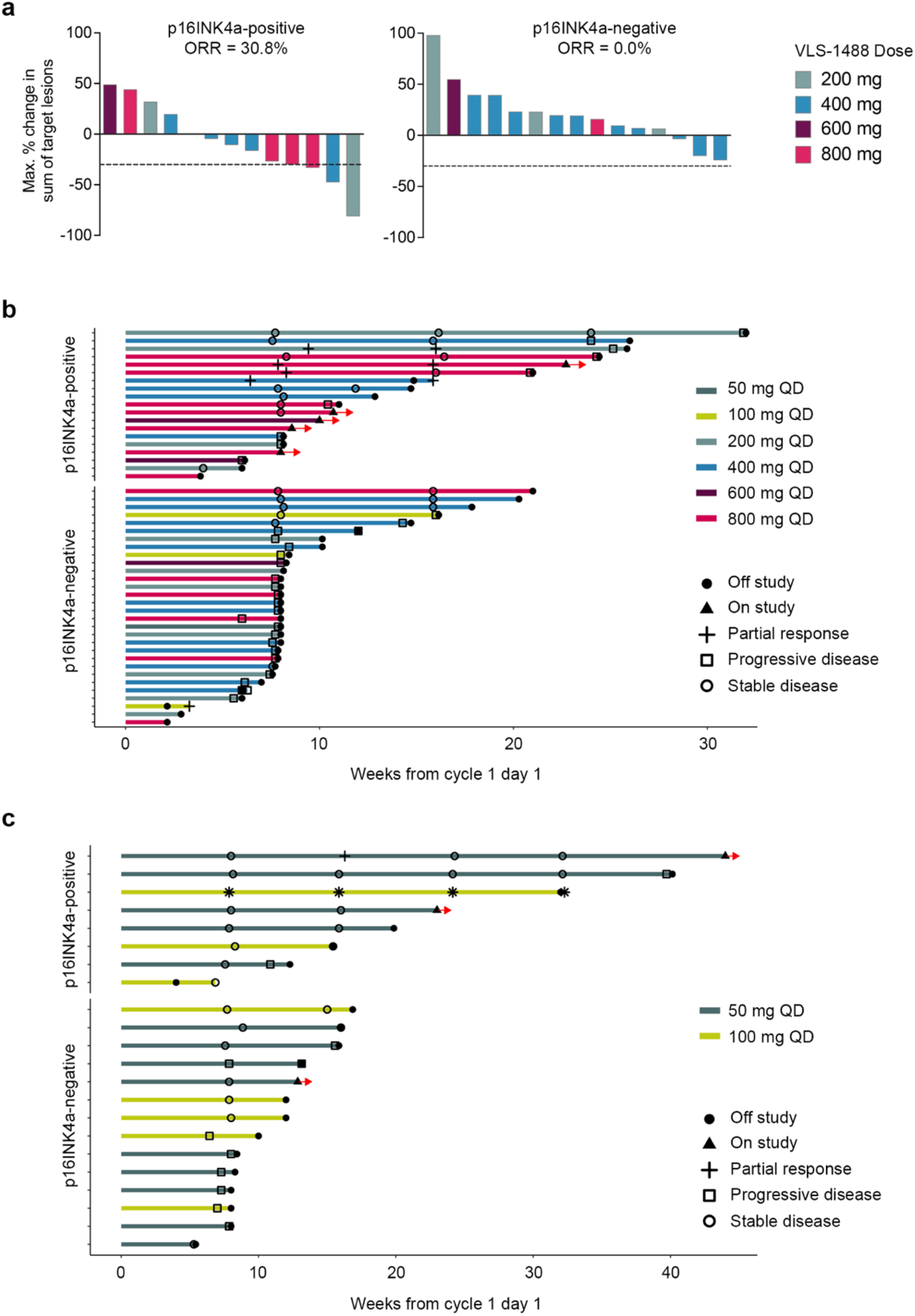
Strong p16INK4a is associated with prolonged duration on treatment with KIF18A inhibitors. **a**, Waterfall plots of maximum percentage change in the sum of target lesions in the response-evaluable, expansion-eligible VLS-1488 subset with 5 or fewer prior lines of therapy and receiving 200-800 mg QD dose, stratified by p16INK4a status and colored by dose level. **b-c**, Swimmer plot for VLS-1488 (b) and sovilnesib (c)-treated patients stratified by p16INK4a status. Bar colors denote dose level. Symbols indicate on-study versus off-study status and best overall response, as shown.

**Supplementary Fig. 7.**
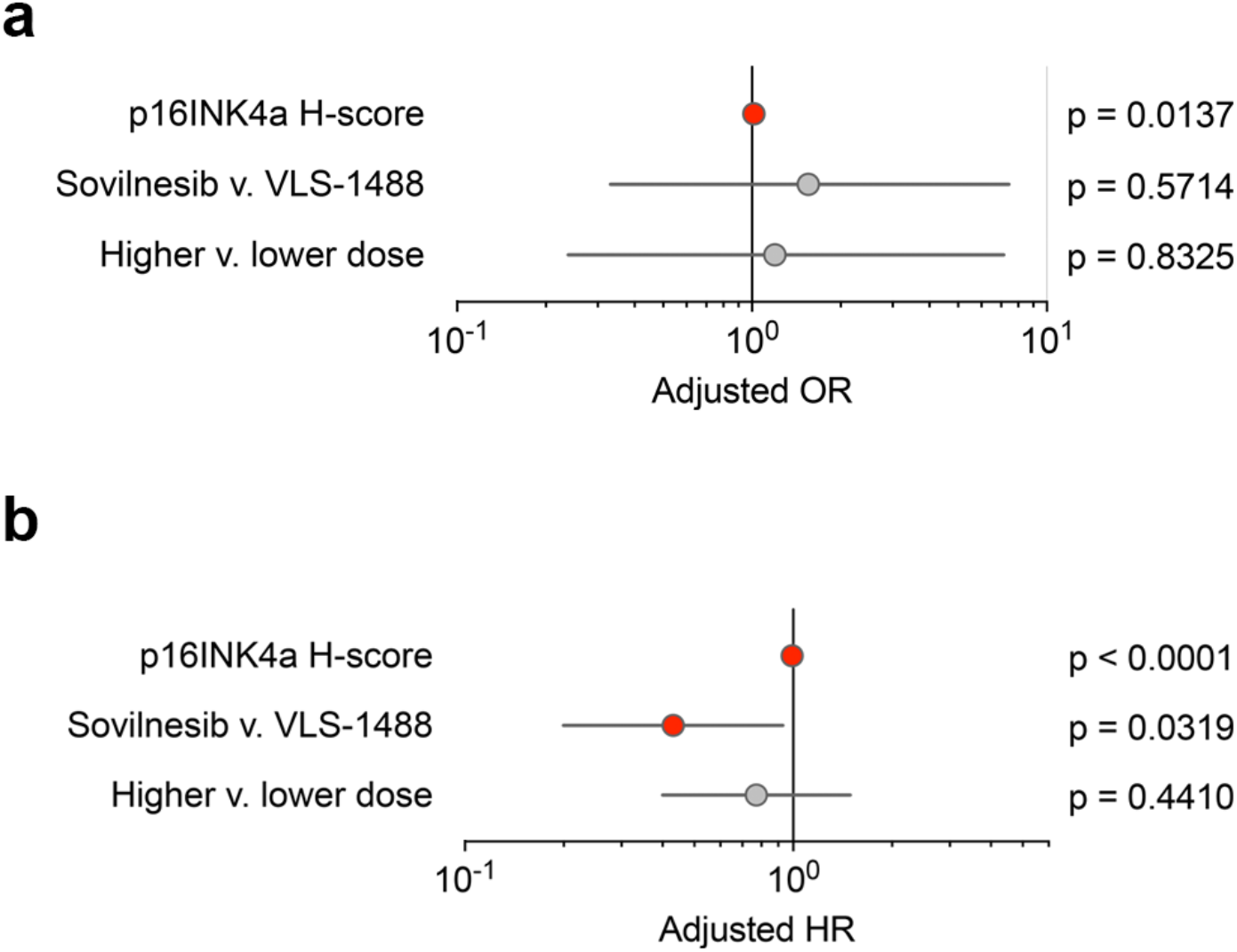
Sensitivity analyses using continuous p16INK4a H-score. **a**, Multivariable logistic regression analysis of objective response in the combined cohort using continuous p16INK4a H-score, with study drug and dose included as covariates. High dose was defined as 100 mg QD or above in sovilnesib and 400 mg QD or above in VLS-1488. **b**, Multivariable Cox regression analysis of progression-free survival in the combined cohort using continuous p16INK4a H-score, with study drug and dose included as covariates. Odds ratios or hazard ratios and 95% confidence intervals are shown.

## TABLES and SUPPLEMENTARY TABLES

**Supplementary Table 1.** Annotation of Rb pathway status in the pan cancer cell line analysis.

| Cell Line | Cancer type | Rb Status | Mechanism | Specific alteration | Source / Pubmed ID | Source identifier |
| --- | --- | --- | --- | --- | --- | --- |
| HCC1187 | Breast Cancer | Altered | CCNE1 / Rb-hyperP | Robinson hyperphosphorylated pRb | PMID | 24265703 |
| HCC1500 | Breast Cancer | Altered | CCNE1 / Rb-hyperP | Robinson hyperphosphorylated pRb | PMID | 24265703 |
| HCC1569 | Breast Cancer | Altered | CCNE1 / Rb-hyperP | Hyperphosphorylated pRb; CCNE1 amplification | PMID | 24265703; 35444283 |
| HCC70 | Breast Cancer | Altered | CCNE1 / Rb-hyperP | Robinson hyperphosphorylated pRb | PMID | 24265703 |
| MDAMB157 | Breast Cancer | Altered | CCNE1 / Rb-hyperP | Robinson hyperphosphorylated pRb | PMID | 24265703 |
| NIHOVCAR3 | Ovarian Cancer | Altered | CCNE1 / Rb-hyperP | CCNE1 amplification | PMID | 35444283 |
| A253 | Head and Neck Cancer | Altered | CDKN2A / p16 lesion | CDKN2A heterozygous frameshift p.Ala17fs*5 | Cellosaurus | CVCL_1060 |
| FADU | Head and Neck Cancer | Altered | CDKN2A / p16 lesion | CDKN2A homozygous splice-site c.151-1G>T | Cellosaurus | CVCL_VP44 |
| HSC3 | Head and Neck Cancer | Altered | CDKN2A / p16 lesion | CDKN2A homozygous truncating mutation p.Glu120Ter | Cellosaurus | CVCL_8323; CVCL_D9WC; CVCL_YA14 |
| LK2 | Lung Cancer | Altered | CDKN2A / p16 lesion | CDKN2A homozygous mutation p.Asp84Tyr (also p.Arg98Leu reported) | Cellosaurus | CVCL_1377 |
| 5637 | Bladder Cancer | Altered | Direct RB1 defect | Cellosaurus homozygous RB1 nonsense | Cellosaurus | CVCL_0126 |
| BT549 | Breast Cancer | Altered | Direct RB1 defect | Cellosaurus homozygous RB1 deletion | Cellosaurus | CVCL_1092 |
| C33A | Cervical Cancer | Altered | Direct RB1 defect | Scheffner 1991 RB mutant/abnormal non-phosphorylated pRB | PMID | 1648218 |
| DU145 | Prostate Cancer | Altered | Direct RB1 defect | Cellosaurus homozygous RB1 nonsense | Cellosaurus | CVCL_0105 |
| DU4475 | Breast Cancer | Altered | Direct RB1 defect | Cellosaurus homozygous RB1 deletion | Cellosaurus | CVCL_1183 |
| HCC15 | Lung Cancer | Altered | Direct RB1 defect | Cellosaurus homozygous RB1 nonsense | Cellosaurus | CVCL_2057 |
| HCC1937 | Breast Cancer | Altered | Direct RB1 defect | Cellosaurus homozygous RB1 in-frame deletion | Cellosaurus | CVCL_0290 |
| MDAMB436 | Breast Cancer | Altered | Direct RB1 defect | Cellosaurus homozygous RB1 frameshift | Cellosaurus | CVCL_0623 |
| MDAMB468 | Breast Cancer | Altered | Direct RB1 defect | Cellosaurus homozygous RB1 deletion | Cellosaurus | CVCL_0419 |
| TCCSUP | Bladder Cancer | Altered | Direct RB1 defect | Oncotarget RB deleted | PMID | 30349650 |
| A549 | Lung Cancer | Intact | Rb-intact | Mizuarai functional RB+ | PMID | 21447152 |
| DLD1 | Colorectal Cancer | Intact | Rb-intact | Mizuarai functional RB+ | PMID | 21447152 |
| HCC1954 | Breast Cancer | Intact | Rb-intact | Robinson intact pRb phosphorylation pattern | PMID | 24265703 |
| HCC38 | Breast Cancer | Intact | Rb-intact | Robinson intact pRb phosphorylation pattern | PMID | 24265703 |
| HCT116 | Colorectal Cancer | Intact | Rb-intact | Mizuarai functional RB+ | PMID | 21447152 |
| HEPG2 | Liver Cancer | Intact | Rb-intact | Mizuarai functional RB+ | PMID | 21447152 |
| HS746T | Gastric Cancer | Intact | Rb-intact | Mizuarai functional RB+ | PMID | 21447152 |
| KATOIII | Gastric Cancer | Intact | Rb-intact | Mizuarai functional RB+ | PMID | 21447152 |
| MDAMB231 | Breast Cancer | Intact | Rb-intact | Robinson intact pRb phosphorylation pattern | PMID | 24265703 |
| NCIH460 | Lung Cancer | Intact | Rb-intact | Mizuarai functional RB+ | PMID | 21447152 |
| NCIN87 | Gastric Cancer | Intact | Rb-intact | Mizuarai functional RB+ | PMID | 21447152 |
| SCC25 | Head and Neck Cancer | Intact | Rb-intact | Mizuarai functional RB+ | PMID | 21447152 |
| SU8686 | Pancreatic Cancer | Intact | Rb-intact | Mizuarai functional RB+ | PMID | 21447152 |
| T24 | Bladder Cancer | Intact | Rb-intact | Mizuarai functional RB+ | PMID | 21447152 |

**Supplementary Table 2.** Cohort composition and evaluable populations for retrospective p16INK4a biomarker analysis across three KIF18A inhibitor cohorts.

|  | Drug | Sovilnesib |  | VLS-1488 |
| --- | --- | --- | --- | --- |
|  | Trial | Amgen<sup>1</sup> | Volastra | Volastra |
| Patients enrolled<sup>2</sup> |  | N/A | 47 | 79 |
| Samples submitted and stained<sup>3</sup> |  | 8 | 29 | 61 |
| Samples excluded from the analysis<sup>4</sup> |  | 0 | 7 | 12 |
| Total Samples Analyzed<sup>5</sup> |  | 8 | 22 | 49 |
| RECIST-evaluable with p16 staining<sup>6</sup> |  | 8 | 21 | 41 |

<sup>1</sup> Only patients with sufficient source data to adjudicate BOR and target lesion measurement were included
<sup>2</sup> Enrolled in the referenced HGSOc cohorts as of the most recent data extraction.
<sup>3</sup> Archival or pre-treatment tumor samples available and processed centrally for p16 IHC as of the most recent data extraction date; patients without available tissue or without consent for pre-treatment biopsy were not submitted.
<sup>4</sup> Excluded for pre-specified QC criteria, including no tissue present on slide, extensive necrosis, or tumor content <20%
<sup>5</sup> Samples passing QC and included in the p16 prevalence analysis.
<sup>6</sup> Subset of analyzed samples with p16 IHC result and RECIST-evaluable clinical outcomes available as of the data extraction. One patient met the RECIST threshold for target lesion shrinkage but was classified as having progressive disease at the first on-treatment assessment and was therefore scored as progressive disease for best overall response.

